# Dual C5aR1 and C5aR2 deficiency uncouples neutrophilic inflammation from antibacterial host defense

**DOI:** 10.64898/2026.11.19.767710

**Authors:** Sarah Walachowski, Archana Jayaraman, Vanessa Tansi, Markus Dudek, Neha Chaudhary, Shuang Xu, Federico Marini, Matthias Gunzer, Christoph Reinhardt, Markus Bosmann

## Abstract

*Streptococcus pneumoniae* is a major cause of bacterial pneumonia and mortality worldwide. While host defense critically depends on complement activation, how C5aR1 and C5aR2 orchestrate pulmonary antibacterial immunity remains unclear. We identified alveolar type II epithelial cells as a local source of C5 and confirmed increased susceptibility of C5-deficient mice to infection. Combined C5aR1/C5aR2 deficiency abolished C5a-induced lung inflammation but paradoxically exacerbated neutrophil accumulation during pneumococcal infection, exceeding that observed in either single-receptor deficiency. Despite their abundance, neutrophils exhibited impaired bacterial clearance, excessive extracellular trap formation, accompanied by aggravated lung injury and mortality. CITE-seq revealed profound neutrophil and macrophage dysregulation, identifying distinct neutrophil subclusters with defective antimicrobial programs, increased cell death and altered inflammatory signatures, consistent with functional exhaustion. Thus, C5aR1 and C5aR2 exert complementary, partially non-redundant functions that couple neutrophil recruitment to antimicrobial competence, prevent excessive inflammation, and preserve effective pulmonary host defense.

## Main

*Streptococcus pneumoniae* (Spn) is a leading bacterial etiology of community-acquired pneumonia in adults^1, 2^, accounting for ∼58 million cases and 634,000 deaths globally in 2023^3^, with disease burden further compounded by the emergence of antibiotic-resistant strains and the limited vaccine effectiveness and coverage^1^. Severe pneumococcal pneumonia can progress to acute respiratory distress syndrome (ARDS), sepsis, and death, underscoring the impact of dysregulated pulmonary inflammation on disease outcomes^1, 4, 5^.

Resident alveolar macrophages (AM) and epithelial cells initiate host defense by sensing invading bacteria and rapidly recruiting neutrophils (PMN) - via chemotactic mediators -, which are critical for pneumococcal clearance and host survival^5, 6^. Upon alveolar infiltration, PMN deploy an antimicrobial arsenal encompassing phagocytosis, cytokine release, and neutrophil extracellular trap (NET) formation^6^. However, PMN activity must be tightly controlled, as excessive or sustained neutrophilic inflammation drives tissue damage and ARDS pathogenesis^4, 7, 8^.

Complement is essential for defense against encapsulated pneumococci^9, 10, 11^, yet overactivation may also contribute to lung injury and ARDS^12^. Complement activation culminates in proteolytic cleavage of C5 into C5a^13^, a potent anaphylatoxin that promotes the recruitment and activation of PMN and AM through the homologous C5aR1 (CD88) and C5aR2 (C5L2)^14,15^. Unlike C5aR1, C5aR2 lacks intracellular G-protein coupling and was initially considered a decoy or scavenger receptor^16, 17, 18^. Subsequent studies, however, have identified active and context-dependent functions for C5aR2 that can either promote ^19, 20, 21^ or restrain inflammation^22, 23, 24^. This complexity is therapeutically as relevant as challenging, since excessive C5a signaling drives inflammation-associated pathology^25, 26, 27, 28^, but complement deficiencies increase susceptibility to invasive pneumococcal disease^29^. Therefore, how C5aR1 and C5aR2 cooperate to coordinate antibacterial defense, while contributing to tissue injury during pneumococcal pneumonia remains unresolved.

Here, using C5aR1/C5aR2 double-deficient mice, we uncover previously unrecognized collaborating functions of C5a receptors in orchestrating pulmonary immunity. Despite complete abrogation of sterile C5a-induced alveolar inflammation, combined receptor deficiency paradoxically exacerbated neutrophilic inflammation, accompanied by PMN dysregulation and exhaustion-like features, and impaired bacterial clearance during pneumococcal pneumonia. These findings identify distinct, partially non-redundant functions of C5aR1 and C5aR2 in coupling neutrophil accumulation to effective antibacterial defense.

## Results

### *De novo* generation and validation of C5aR1/2^-/-^ double-deficient mice

Mechanistic dissection of C5a signaling has been limited by the lack of combined C5aR1/C5aR2 deletion, as the genomic proximity of both receptors precludes conventional intercrossing of existing single-knockout strains. To overcome this constraint, we engineered deletion of the C5aR1/C5aR2 locus by CRISPR–Cas9-mediated germline editing. Single-guide RNAs targeting exon 2 of the *C5ar1* gene and exons 1-2 of the *C5ar2* gene on chromosome 7 were microinjected into C57BL/6J zygotes (**Fig. 1a**). Sanger sequencing of F0 founders confirmed a 12.6 kb deletion encompassing the coding regions of both genes. PCR-based genotyping identified an 844 bp mutant amplicon specific to the recombined allele (**Fig. 1b**). Dual C5aR1/C5aR2 deficiency was confirmed *in vivo*, as alveolar macrophages (AM) and neutrophils (PMN) from C5a-stimulated or sham-treated C5aR1/2^-/-^ mice lacked detectable receptor expression (**Fig. 1c-e**). C5aR1/2^-/-^ mice were viable and fertile, with a normal sex distribution and litter frequency, although litter sizes were modestly reduced in homozygous breeding schemes (**Extended Data Fig. 1a-c**). Body and organ weights were comparable to wildtype (WT) controls, as were plasma and serum biochemical parameters including proteins, metabolites, electrolytes and lipids (**Extended Data Fig. 1d-i**). Circulating leukocyte, erythrocyte, and platelet counts were normal in C5aR1/2^-/-^ mice (**Extended Data Fig. 1j-k**). Consistently, splenic immunophenotyping of naïve male and female C5aR1/2^-/-^ mice showed no relevant alterations in immune cell populations (**Extended Data Fig. 2**). Together, these data establish C5aR1/2^-/-^ mice as a robust model for studying C5a receptor biology without detectable defects in basal immune development or systemic homeostasis.

**FIGURE 1.**
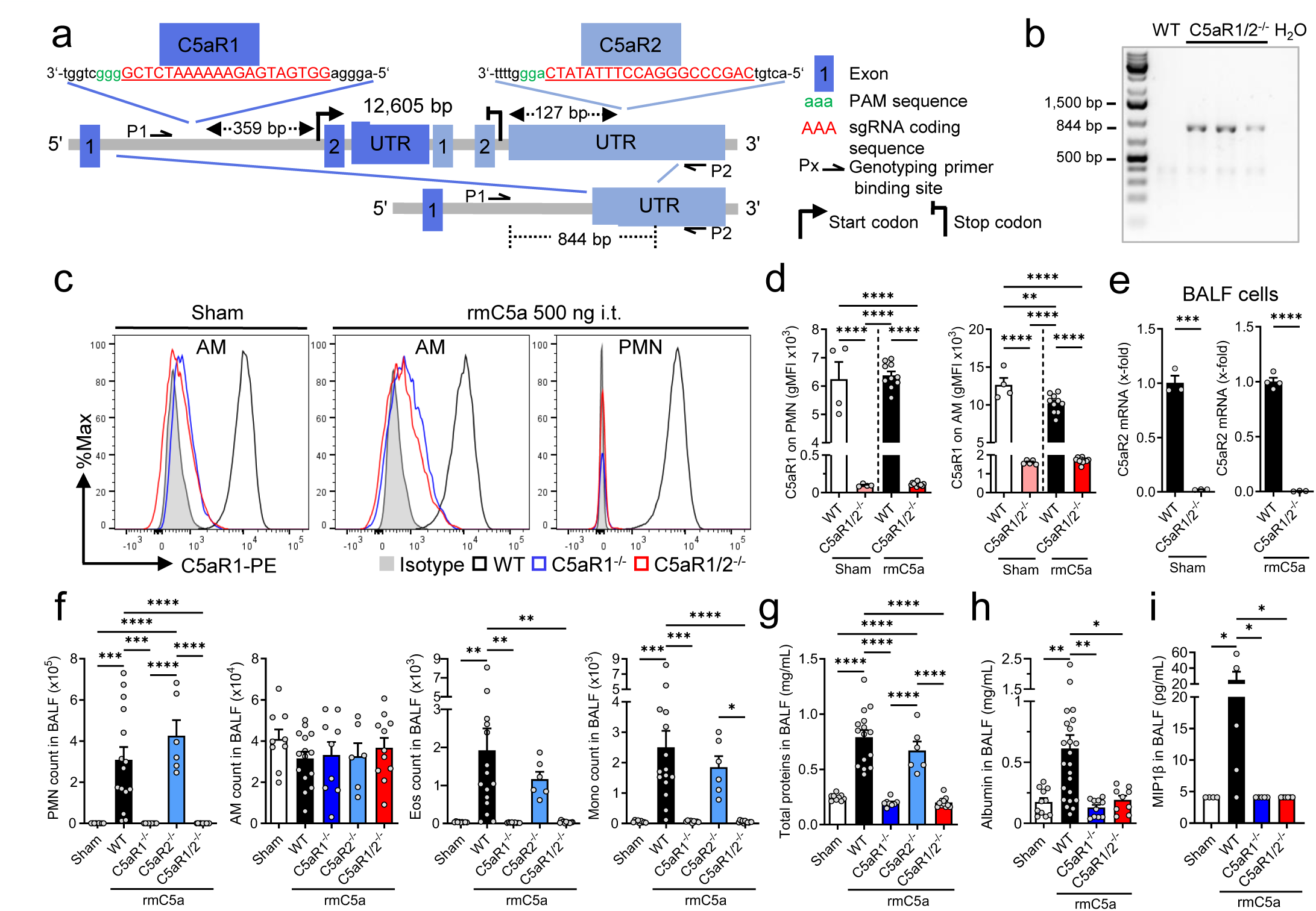
Generation and characterization of C5aR1 and C5aR2 double knock-out mice. (**a-b**) C5aR1/2^-/-^ double knock-out mice were generated by a homologous CRISPR-Cas9 induced 12.6 kb deletion of the coding regions of C5aR1 (exon 2) and C5aR2 (exons 1 and 2) on chromosome 7 in C57BL/6J (WT) mice. (**a**) Scheme of C5aR1 and C5aR2 loci targeting. (**b**) Representative gel of the mutant 844 bp PCR product exclusively present in homozygous C5aR1/2^-/-^ mice. Lane 1: DNA Ladder, Lane 2: WT control DNA deprived of the mutant band, Lanes 3-5: DNA samples of n = 3 double knock-out mice showing the 844 bp mutant band, Lane 6: PCR negative control (H2O). A faint nonspecific band of ∼350 bp was observed in both genotypes. (**c-i**) *In vivo* analysis of alveolar inflammatory response in PBS (sham) or rmC5a (500 ng) intratracheally (i.t.) instilled WT, C5aR1^-/-^, C5aR2^-/-^ and C5aR1/2^-/-^ mice after 7 h. (**c**) Representative histograms and (**d**) bar graphs of cell surface C5aR1 staining on Ly6G^+^Ly6C^+^ neutrophils (PMN) and SiglecF^+^CD11c^+^ alveolar macrophages (AM); flow cytometry (sham, n = 4 mice/group; rmC5a, n = 10-11 mice/group). (**e**) C5aR2 mRNA in BALF cells isolated from sham or rmC5a-treated WT and C5aR1/2^-/-^ mice (n = 3-4 mice/group); RT-qPCR. (**f**) Live AM, PMN, Ly6G^-^Ly6C^+^ monocytes (Mono) and SiglecF^+^CD11c^-^ eosinophils (Eos) in BALF; flow cytometry (n = 6-15 mice/group). Quantification of (**g**) total protein (n = 6-15 mice/group), (**h**) albumin (n = 9-23 mice/group) and (**i**) MIP-1β (n = 4-6 mice/group) by BCA assay or ELISA in BAL. Data are combined (**d**, **f-i**) or representative (**b, c, e**) of 2-3 independent experiments. Symbols represent one individual mouse, and bars indicate mean ± SEM. *p < 0.05, **p < 0.01, ***p < 0.001, ****p < 0.0001, Student’s t-test analysis or one-way ANOVA. UTR, untranslated region; PAM, protospacer adjacent motif; gMFI, geometric mean of fluorescence; rmC5a, recombinant mouse C5a; BALF, bronchoalveolar lavage fluid.

### Dual C5aR1/2 deletion confers resistance to C5a-induced acute lung injury

To determine whether combined C5aR1 and C5aR2 deficiency suppresses C5a-driven inflammation *in vivo*, we used an established model of C5a-induced acute lung injury (C5a-ALI)^30^. C5a-induced influx of neutrophils, eosinophils and monocytes was completely abrogated in C5aR1/2^-/-^ mice, yet AM counts remained comparable between both genotypes and were unaffected by C5a-ALI (**Fig. 1f**, **Extended Data Fig. 3a-c**). C5a promoted alveolar-capillary barrier disruption, and induced chemokines (MIP-1β) in WT but not C5aR1/2^-/-^ mice with elevated total protein and albumin concentrations in BALF (**Fig. 1g-i**). Deletion of C5aR1 alone recapitulated the phenotype of C5aR1/2^-/-^ mice, completely abolishing myeloid influx and alveolar injury, whereas resistance to C5a-ALI was limited in C5aR2^-/-^ mice (**Fig. 1f-i**, **Extended Data Fig. 3a-c**), thus identifying C5aR1 as the dominant receptor driving C5a-ALI.

### C5a reprograms alveolar macrophages during acute lung injury

To further elucidate the effects of C5a on AM, we performed transcriptomics. Bulk RNA sequencing of FACS-sorted live SiglecF^+^CD11c^+^CD64^+^ AM from BALF of sham and C5a-ALI mice showed marked C5a-driven transcriptional reprogramming, with 822 upregulated and 957 downregulated genes in C5a-exposed AM relative to controls (**Fig. 2a-b**). KEGG analysis identified enrichment of ribosomal, lysosomal, peroxisomal, apoptotic, ferroptotic, and C-type lectin receptor and PPAR pathways, whereas Gene Ontology (GO) analysis revealed programs associated with inflammation, cell proliferation, cytokine responses, signal transduction, cell death, and homeostasis (**Fig. 2c-d**). Curated heatmaps and network analysis indicated coordinated activation of myeloid, chemotactic, antibacterial and stress-response programs, with increased expression of *Hmox1*, *Cd274*, *Ripk1*, *CD200r1* and *Hif1a,* and reduced expression of genes linked to adhesion, tissue repair and homeostasis, including *Itgb1*, *Tgfb1*, *Itgal* and *Sirpa* (**Fig. 2e-g** and **Extended Data Fig. 3d-e**). Differentially expressed genes further reflected enhanced cell death (*Bax*, *Ripk1*), myeloid recruitment (*Ccl2*, *Ccl7*), inflammatory signaling (*S100a8*, *Tbk1*), myeloid activation (*Cd63*, *Cd84*), immune regulation (*Cd200r1*, *Cd274, Cd101*), combined with compromised macrophage differentiation (*Myadm*), tissue repair (*Tgfb1*) and phagocytosis (*Sirpa*) (**Fig. 2h**). Together, these findings position C5a as a key instructive signal driving AM reprogramming in acute lung injury.

**FIGURE 2.**
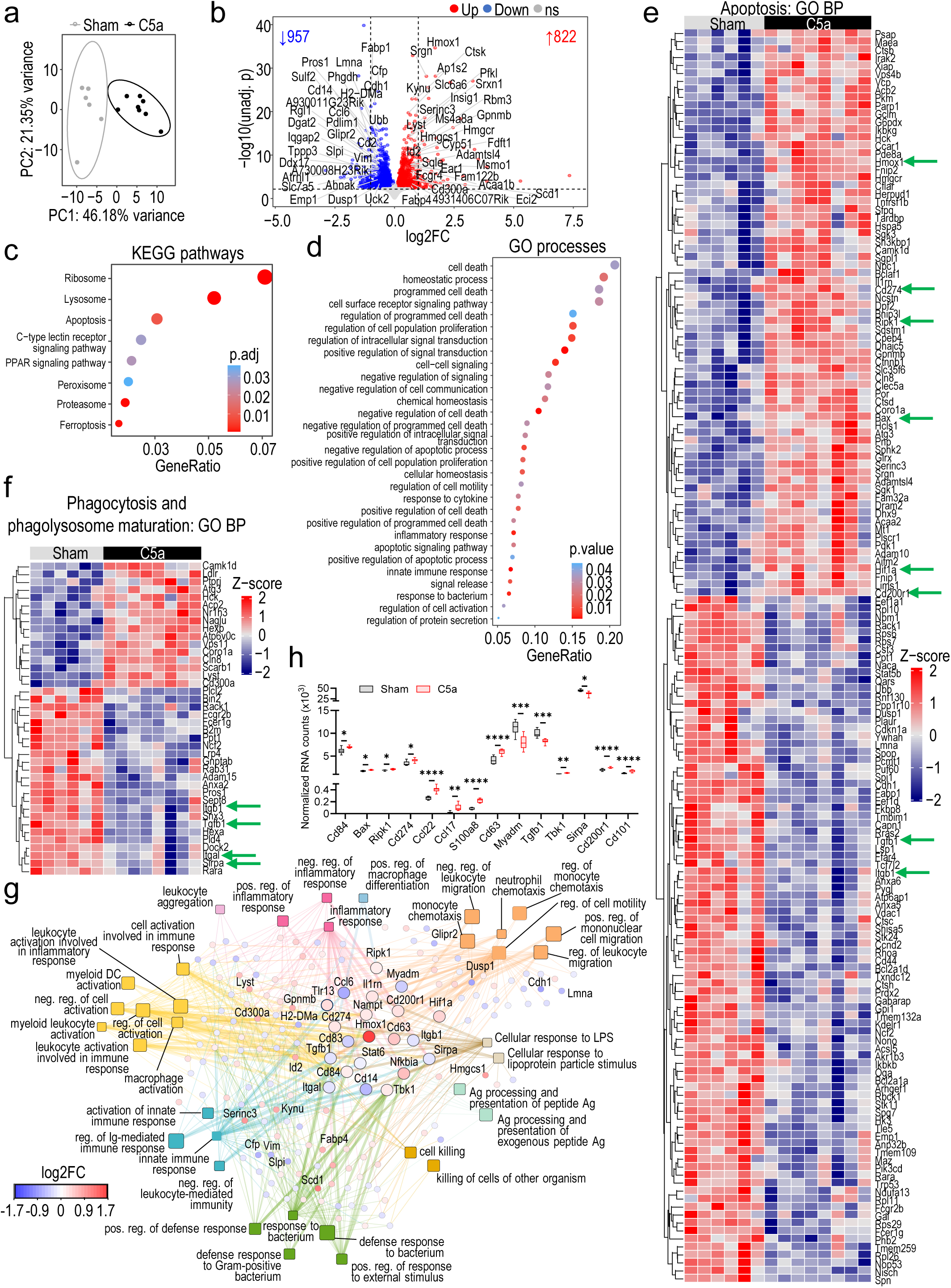
C5a alters alveolar macrophages transcriptional programs governing inflammation, phagocytosis and apoptosis. Bulk RNA-seq of FACS-sorted live SigF^+^CD11c^+^CD64^+^ AM from sham mice and C5a-induced ALI mice (rmC5a, 500 ng i.t., 7 h; n = 6-8 mice/group). (**a**) Principal component analysis (PCA) of the top 500 most variable genes after variance stabilizing transformation of RNA-seq counts from control (sham) and C5a-ALI mice. (**b**) Volcano plot showing differentially expressed genes (DEGs) in C5a-ALI vs. sham mice. The number of significant upregulated and downregulated genes is indicated. The horizontal dotted line represents adjusted p (adj.p) = 0.05 and the vertical dotted lines represent |log2 fold change| = 1. Significant DEGs with baseMean > 1000 are labelled. FC, Fold Change; unadj. p, unadjusted p; ns, not significant. (**c**) Dot plot showing selected pathways from KEGG enrichment analysis. (**d**) Dot plot showing selected Gene Ontology (GO) Biological Processes (BPs) relevant to macrophage immune response. (**e-f**) Heatmaps showing significant DEGs enriched in GO BPs associated with (**e**) apoptosis and cell death, or (**f**) phagocytosis and phagolysosomal maturation. Heatmaps show normalized expression values of DEGs with baseMean ≥1000 (Z-score from -2, blue to +2, red). Genes of interest are indicated with a green arrow. (**g**) Network plot of DEGs with baseMean ≥1000 enriched in GO BPs associated with myeloid cell activation, and leukocyte migration and chemotaxis. Squares represent GO BPs and circles represent DEGs. Relevant DEGs or those with |log2FC| > 0.58 are labelled. GO BPs were manually grouped and colored according to functional similarity. DEG color indicates log2FC (-1.7, blue to +1.7, red). Relevant DEGs are shown as larger circles for visualization. Edge color reflects the corresponding GO BP, and thicker edges indicate the links to relevant DEGs. Square size is proportional to the enrichment p.value elim. (**h**) Box plot of normalized counts for selected immune-related genes in sham and C5a-ALI mice. See also **Extended Data** Fig. 3.

### Alveolar type 2 epithelial cells are a local source of C5 in the lung

Previous studies, including single-cell RNA-seq analyses from our laboratory, identified C5 transcripts in alveolar type 2 (AT2) epithelial cells in mouse and human lungs^31^, suggesting local pulmonary C5 expression. To evaluate C5 production, we used naive C5^lacZ^ reporter mice, in which β-galactosidase (β-gal) activity serves as a surrogate for C5 expression. X-gal staining revealed robust blue cytoplasmic reporter activity in lung cells (**Fig. 3a**), in addition to the expected hepatic signal (**Extended Data Fig. 4a**), with much lower expression in other organs (**Extended Data Fig. 4b**). Co-staining for C5 and CD45 showed no colocalization, consistent with non-hematopoietic cells being the predominant local source of pulmonary C5 (**Fig. 3b**). To assess for epithelial expression, we used the β-galactosidase substrate (DDAOG), which is enzymatically hydrolyzed to far-red fluorescent DDAO^32^. Confocal microscopy revealed prominent cytoplasmic DDAO fluorescence in surfactant protein C (SPC)^+^ AT2 cells of naive C5^lacZ^ mice, but not WT lungs (**Fig. 3c**). For independent validation, C5^LacZ^ reporter activity was confirmed in isolated AT2 cells by flow cytometry (**Fig. 3d**, **Extended Data Fig. 4c**). *Ex vivo* IVIS imaging of intact lung lobes corroborated prominent pulmonary C5 expression (**Fig. 3e**). In conclusion, these findings establish AT2 cells as the principal local source of complement C5, establishing a reservoir for C5a generation within the alveolar niche.

**FIGURE 3.**
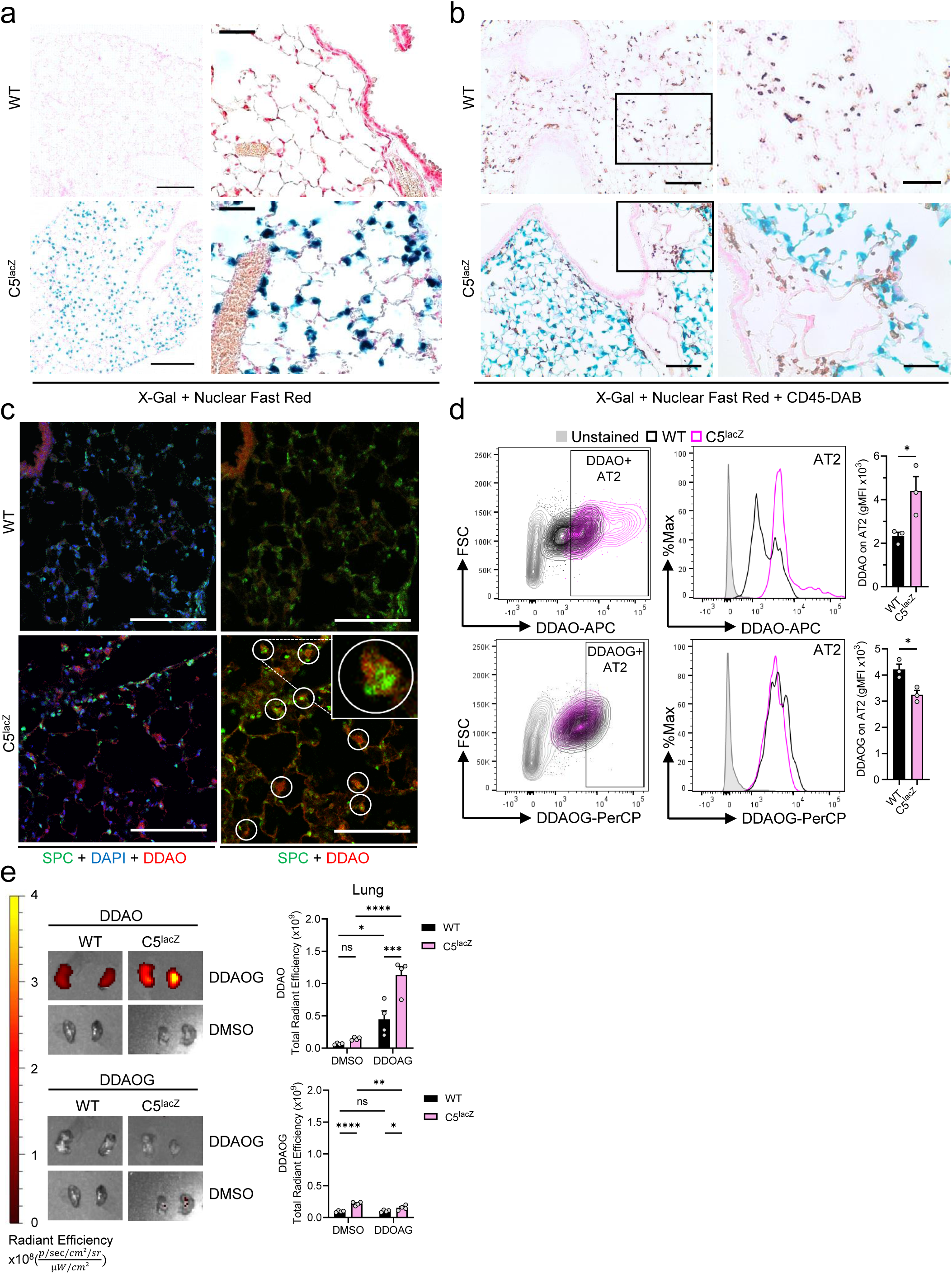
C5 is locally produced in alveolar type 2 cells (AT2). Lungs from naïve C5^lacZ^ and WT control mice were collected for immunostaining and flow cytometry analysis. (**a-b**) Representative images of immunohistochemistry analysis for X-Gal and Nuclear Fast Red staining of lung sections. (**a,** left panels) Fixed-frozen sections, scale bars = 200 μm. (**a**, right panels) Paraffin-embedded sections, scale bars = 50 μm. (**b**) Paraffin-embedded lung sections were co-stained for CD45. Boxed regions were magnified and presented on the right side of the respective panels. Scale bars = 100 μm (whole sections) and 50 μm (zoomed sections). (**c**) Representative fixed-frozen lung sections analyzed by immunofluorescence for Surfactant Protein C (SPC, green), the hydrolyzed product DDAO (red) derived from DDAOG substrate to monitor C5^lacZ^ reporter, and DAPI (blue). White circles show co-localization of DDAO and SPC staining within the same cells but with distinct subcellular localization. Scale bars = 100 μm. (**a-c**) Data are representative of 3 experiments. (**d**) Representative overlay contour plots, histograms, and bar graphs of geometric mean fluorescence intensities (gMFI) of DDAOG and its cleavage product, DDAO in CD45^-^ CD326^+^CD24^-^Pdpn^-^MHCII^+^ AT2 cells; flow cytometry. Representative gating strategy is presented in Extended Data Fig. 4c. (**e**) *Ex vivo* IVIS images and reporter activity of lung lobes isolated from C5^lacZ^ and WT mice within minutes after DDAOG substrate or DMSO (control) injection. Data are representative of 2 independent experiments. Symbols represent one individual mouse (n = 3 mice/group), and bars indicate mean ± SEM. *p < 0.05, **p < 0.01, ***p < 0.001, ****p < 0.0001, ns, not significant, as determined by (**d**) Student’s t-test analysis or (**e**) one-way ANOVA. Pdpn, podoplanin.

### C5-deficient mice are highly susceptible to streptococcal pneumonia

To assess the protective role of C5, we challenged C5-deficient mice with the *Streptococcus pneumoniae* TIGR4 strain (Spn4), an invasive clinical isolate^33^. C5 deficiency drastically reduced 14-day survival rates (**Fig. 4a**), and exacerbated alveolar-capillary barrier disruption, as indicated by elevated BALF total protein concentrations at 18 h post-infection (**Fig. 4b**). Consistently, alveolar PMN numbers were higher in C5-deficient mice compared with WT mice (**Fig. 4c-e**). In contrast, monocytes and AM numbers and percentages were significantly reduced, whereas eosinophil counts were similar… Notably, C5aR1 expression on PMN and AM was moderately higher in C5-deficient mice (**Fig. 4f-g**). Collectively, these findings identify C5, the substrate for C5a generation, as a critical determinant of host protection during pneumococcal pneumonia, limiting pulmonary inflammation and tissue injury while supporting antimicrobial defense.

**FIGURE 4.**
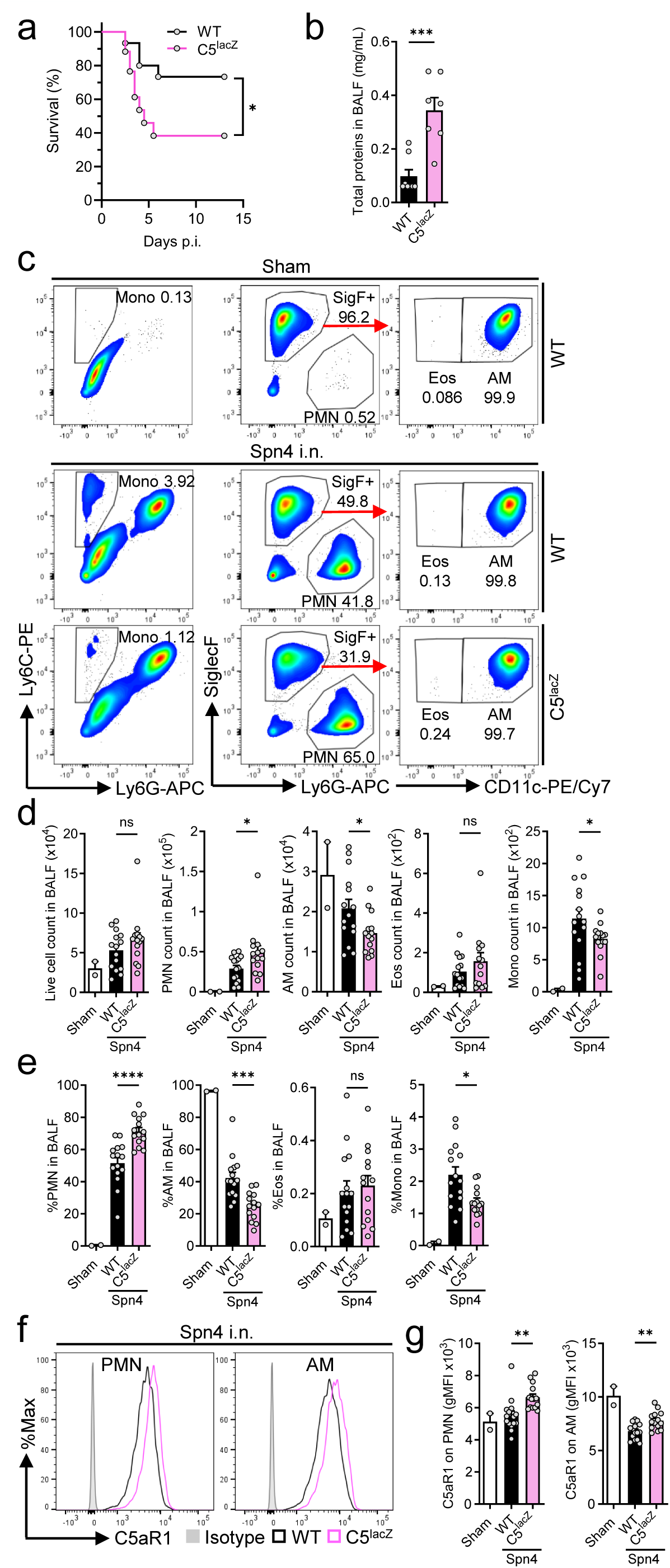
Complement component C5 coordinates host defense during pneumococcal infection. (**a-g**) C5^lacZ^ and WT control mice were intranasally (i.n.) challenged with *S. pneumoniae* (1-2x10^6^ CFU) or PBS (sham), and BALF were collected 18 h later. (**a**) Survival was monitored for 14 days (n = 15-17/group). (**b**) Total protein concentration in BALF supernatants was measured using a Bradford assay (n = 7/group). (**c**) Representative flow cytometry plots of Ly6G^-^Ly6C^+^ monocytes (Mono), Ly6G^+^ neutrophils (PMN), SiglecF^+^CD11c^+^ alveolar macrophages (AM) and SiglecF^+^CD11c^-^ eosinophils (Eos) in BAL. Cell frequencies of live populations are indicated next to each gate. (**d**) Absolute numbers and (**e**) percentages of live BALF were determined by flow cytometry (n ≥ 15/group). (**f**) C5aR1 surface expression on each BALF cell population expressed in gMFI (n ≥ 15/group). (**g**) Representative histograms of C5aR1 expression on PMN and AM. Isotype control (grey-filled histogram) was included in each panel. Data are shown as mean ± SEM and are representative of 2-3 independent experiments. Each circle represents an individual mouse. *p < 0.05, **p < 0.01, ***p < 0.001, ****p < 0.0001, ns, not significant, as determined by one-way ANOVA. Kaplan-Meier survival curve was analyzed by Log-rank test. rmC5a, recombinant mouse C5a; BALF, bronchoalveolar lavage fluid; FMO, fluorescence minus one. SigF, SiglecF.

### Combined C5aR1-C5aR2 deficiency exacerbates pneumococcal pneumonia and impairs antibacterial host defense

We next investigated whether the phenotype observed in C5-deficient mice infected with Spn4 extends to deficiency of both C5a receptors. Strikingly, alveolar neutrophil accumulation was increased in C5aR1^-/-^ mice and further exacerbated in C5aR1/2^-/-^ mice compared to WT mice, while C5aR2^-/-^ mice also showed difference trend towards higher PMN numbers (**Fig. 5a-c**). Conversely, AM abundance was more markedly reduced in C5aR1/2^-/-^ mice than in the other genotypes. Eosinophil and monocyte counts were also more elevated in C5aR1/2^-/-^ mice (**Fig. 5a-c**). Combined receptor deficiency led to aggravated lung injury, as indicated by increased BALF albumin and total protein (**Fig. 5d-e**), accompanied by increased concentrations of cytokines/chemokines, including IL-6, TNF-α, KC/CXCL1, IL-1β, IFNƴ, CXCL10 and CXCL5 (**Fig. 5f-h**).

**FIGURE 5.**
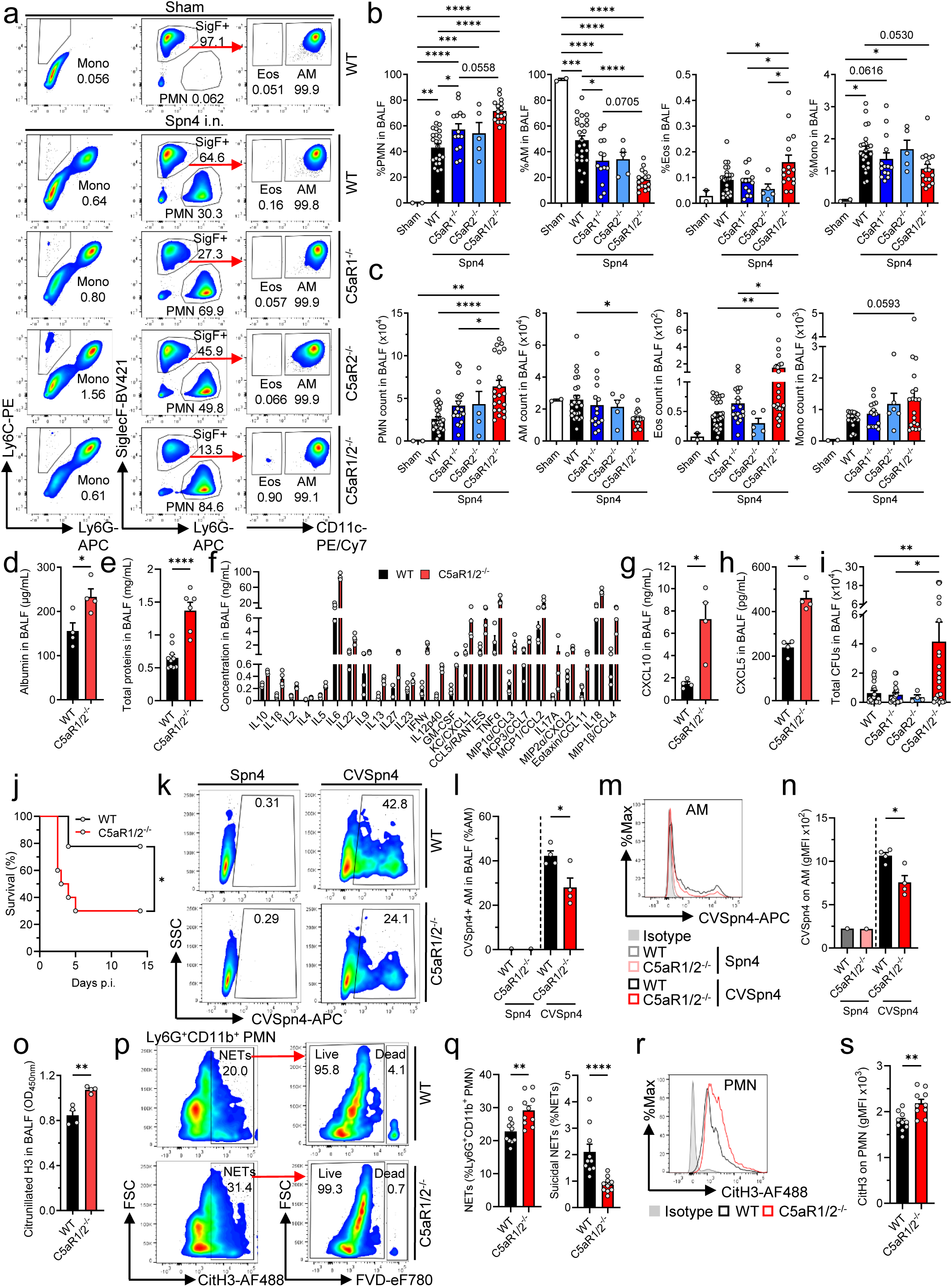
Dual C5aR1 and C5aR2 deficiency exacerbates pneumococcal infection. (**a-j**, **o-s**) WT, C5aR1^-/-^, C5aR2^-/-^ and C5aR1/2^-/-^ mice were intranasally (i.n.) infected with *S. pneumoniae* (1-2x10^6^ CFU) or treated with PBS (sham), and BALF were collected after 18 h. (**a**) Representative flow cytometry plots of Ly6G^-^Ly6C^+^ monocytes (Mono), Ly6G^+^ neutrophils (PMN), SiglecF^+^CD11c^+^ alveolar macrophages (AM) and SiglecF^+^CD11c^-^ eosinophils (Eos) in BAL. Cell frequencies of live populations are indicated next to each gate. (**b**) Percentages and (**c**) absolute numbers of live BALF were determined by flow cytometry (n ≥ 15/group except for C5aR2^-/-^ mice group where n = 5). (**d**) Albumin and (**e**) total protein concentration in BALF supernatants were measured using a Bradford assay or ELISA, respectively (n ≥ 4/group). (**f**) Concentration of 26 cytokines and chemokines were measured in BALF supernatants by multiplex immunoassay. Analytes that showed significant differences between groups are displayed (n = 4/group). (**g**) CXCL10 and (**g**) CXCL5 were measured in BALF supernatants by ELISA (n = 4/group). (**i**) CFUs in BALF were quantified (n ≥ 20/group). (**j**) Survival was monitored for 14 days (n = 9-10/group). (**k-n**) WT and C5aR1/2^-/-^ mice were intranasally (i.n.) infected with CellVue-labelled *S. pneumoniae* (CVSpn4), unlabeled *S. pneumoniae* or treated with PBS (sham) for 90 min, and BALF were collected for flow cytometry analysis (n = 4 mice/group). (**k**) Representative flow cytometry plots and (**l**) Percentage of alveolar macrophages (AM) associated with fluorescent bacteria. (**m**) Representative histogram and, (**n**) geometric mean fluorescence intensity (gMFI) of *S. pneumoniae* uptake by AM. (**o**) Citrullinated histone 3 (CitH3) as a marker for NETs in BALF supernatants (ELISA, n = 4/group). (**p**) Representative flow cytometry analysis for CitH3 on Ly6G^+^CD11b^+^ PMN and (**q**) fraction of PMN undergoing NETosis expressed as the percentage of total CitH3^+^ PMN. The fraction of suicidal/lytic NETs expressed as the percentage of dead CitH3^+^ PMN is also shown (n = 10/group). (**r**) Representative overlay histograms and (**s**) gMFI of CitH3 on PMN. Data are shown as mean ± SEM and are representative of 2-3 independent experiments. Each circle represents an individual mouse. *p < 0.05, **p < 0.01, ***p < 0.001, ****p < 0.0001, as determined by Student’s t-test or one-way ANOVA when more than 2 groups were compared. Kaplan-Meier survival curve was analyzed by Log-rank test. BALF, bronchoalveolar lavage fluid. FVD, Fixable Viability Dye; SigF, SiglecF.

Despite enhanced PMN influx, C5aR1/2^-/-^ mice exhibited higher alveolar bacterial burden and increased mortality following infection, suggesting that the amplified inflammatory response did not translate into improved bacterial clearance (**Fig. 5i-j**). To determine whether impaired phagocytic function contributed to this defect, we quantified binding and uptake of fluorescently labeled Spn4 by AM *in vivo*. At 1.5 hours post-infection, ∼42% of WT AM were bacteria-positive compared with ∼24% in C5aR1/2^-/-^ mice (**Fig. 5k-l**). Bacteria-positive C5aR1/2^-/-^ AM also displayed lower fluorescence intensities, indicative of a reduced relative bacterial association per cell, consistent with impaired phagocytic capacity (**Fig. 5m-n**).

Given the involvement of C5a in NET formation^34^, we next assessed NETosis during pneumococcal pneumonia. Citrullinated H3 (CitH3), a marker of NETs, was increased in BALF from Spn4-infected C5aR1/2^-/-^ mice and correlated with inflammatory and lung injury markers (**Fig. 5o**). Flow cytometry confirmed enhanced NET formation in C5aR1/2^-/-^ neutrophils, higher frequencies of CitH3^+^ PMN and greater CitH3 fluorescence intensity (**Fig. 5p-s**). NET formation occurred predominantly through non-lytic (‘vital’) NETosis in both genotypes, although lytic (‘suicidal’) NETosis was lower in C5aR1/2^-/-^ PMN (**Fig. 5p-q**). Elevated NETs in C5aR1/2^-/-^ mice coincided with increased alveolar-capillary barrier permeability, consistent with their contribution to lung injury^35^.

Collectively, these findings indicate that intact and coordinated C5aR1-C5aR2 signaling limits neutrophilic inflammation while supporting phagocyte-mediated bacterial clearance during pneumococcal pneumonia.

### C5aR1 and C5aR2 shape alveolar macrophage transcriptional responses during pneumococcal infection

To define macrophage-intrinsic inflammatory programs and the functional role distributions of the two C5a receptors during bacterial pneumonia, we performed bulk RNA sequencing of AM isolated from WT, C5aR1^-/-^, C5aR2^-/-^ and C5aR1/2^-/-^ mice after Spn4 infection. Principal component analysis (PCA) revealed genotype-dependent clustering of AM transcriptional profiles, reflecting extensive reprogramming in the absence of C5aR signaling (**Fig. 6a**). AM lacking C5aR1, either alone or combined with C5aR2 deficiency, showed the greatest divergence from WT cells, whereas C5aR2^-/-^ AM partially overlapped with WT and, to a lesser extent, C5aR1/2^-/-^, indicating a less pronounced transcriptional shift (**Fig. 6a**). Differential expression analysis identified shared and genotype-specific signatures across AM genotypes (**Fig. 6b-c**). C5aR1/2^-/-^ AM displayed the largest number of differentially expressed genes (DEGs) relative to WT cells, while changes in C5aR2^-/-^ AM were more limited (**Fig. 6b, d**). Genes associated with innate immune and inflammatory pathways (including cytokine, interleukin, interferon and TLR signaling, and autophagy) showed distinct expression patterns across genotypes (**Fig. 6c**), suggesting divergent and non-redundant roles for C5aR1 and C5aR2. A subset of genes was strongly upregulated in C5aR1/2^-/-^ AM (**Fig. 6c-d**), indicative of an enhanced inflammatory response.

**FIGURE 6.**
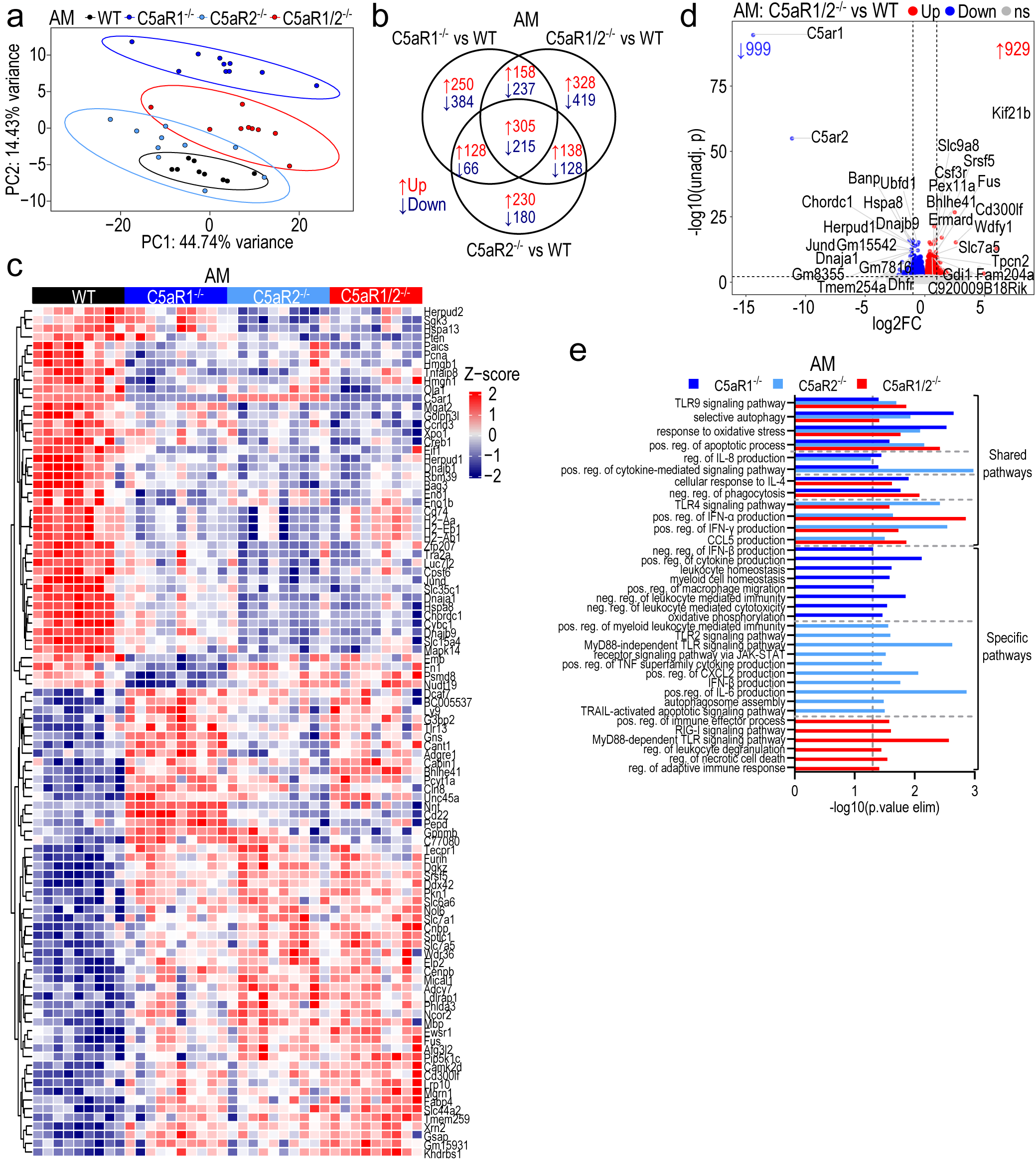
Loss of C5aR1 and C5aR2 disrupts macrophage transcriptional programs during *S. pneumoniae* infection. Bulk RNA-seq was performed on FACS-sorted live SiglecF^+^CD11c^+^CD64^+^ alveolar macrophages (AM) from WT, C5aR1^-/-^, C5aR2^-/-^ and C5aR1/2^-/-^ mice 18 h after intranasal infection with 2x10^6^ CFU *S. pneumoniae*. All comparisons are shown relative to WT controls. (**a**) Principal component analysis (PCA) of the top 500 most variable genes across samples (n = 9-10 mice/group). (**b**) Venn diagram of shared and genotype-specific significant differentially expressed genes (DEGs). (**c**) Heatmap of normalized expression values of top 50 significant DEGs (baseMean ≥ 1000; Z-score from -2, blue to +2, red). (**d**) Volcano plot showing DEGs in C5aR1/2^-/-^ vs. WT AM. The number of significantly upregulated and downregulated genes is indicated. The horizontal dotted line represents adjusted p (adj. p) = 0.05 and the vertical lines represent |log2 fold change| = 1. Top 15 significant up-/downregulated DEGs with |log2FC| > 0.58 and baseMean > 100 are labelled. FC, Fold change; unadj. p, unadjusted p; ns, not significant. (**e**) Gene Ontology (GO) biological processes enriched among significant DEGs, highlighting immune-related pathways. pos., positive; neg., negative; reg., regulation.

GO analysis of significant DEGs identified shared enrichment of processes and genotype-specific programs. A core set of immune processes, including TLR9 signaling, autophagy, oxidative stress response, apoptosis, cytokine signaling, phagocytosis and IFNα/ƴ production, was significantly overrepresented across several or all three C5aR-deficient genotypes, reflecting a canonical activation program during infection (**Fig. 6e**), although distinct genotype-specific enrichments also emerged. C5aR1 deficiency favored regulatory and homeostatic responses, including negative regulation of cytokine/IFNα production and leukocyte-mediated immunity, and leukocyte homeostasis, whereas C5aR2 deficiency was associated with immune effector pathways positive regulation, including TLR2- and MyD88-independent signaling, cytokine production (TNFα, IFNβ, IL6 and CXCL2) and apoptosis. Conversely, C5aR1/2^-/-^ AM enriched MyD88-dependent signaling, leukocyte degranulation, cell death and RIG-I signaling processes.

Gene set enrichment analysis (GSEA) further resolved coordinated pathway changes (**Extended Data Fig. 5**). C5aR1^-/-^ AM showed enrichment of TGFβ signaling pathway and reduced oxidative phosphorylation, whereas C5aR2^-/-^ AM exhibited repression of multiple inflammatory hallmark gene sets, including apoptosis, complement, TNFα and IFNα/ƴ responses. By contrast, IL6-JAK-STAT3 and IFNα/ƴ signaling were enhanced in C5aR1/2^-/-^, reflecting hyperinflammation. Thus, dual C5aR1/2 deficiency promoted a transcriptional state not recapitulated by either single knockout, suggesting independent role for the two receptors in macrophage activation.

We next examined similar changes that occurred in PMN. In contrast to AM, PMN exhibited only moderate genotype-dependent transcriptional reprogramming, with C5aR1/2^-/-^ PMN largely overlapping with WT and displaying fewer DEGs than either single knock-out populations (**Extended Data Fig. 6a-d**). GO analysis revealed selective modulation of immune and regulatory processes in C5aR1/2^-/-^ PMN, including ERK1/2 signaling, apoptotic process, cellular response to IL7 and LIF, regulation of type I IFN, IL10, neutrophil migration and TGFβ signaling (**Extended Data Fig. 6e**). GSEA showed enrichment of oxidative phosphorylation and ROS pathways without consistent increase in canonical inflammatory signatures (**Extended Data Fig. 6f**).

In conclusion, these findings reveal distinct and interdependent contributions of C5aR1 and C5aR2 to myeloid transcriptional responses during pneumococcal infection with combined receptor loss preferentially amplifying inflammatory programs in AM.

### CITE-seq reveals dysregulated alveolar myeloid states during pneumococcal infection

To investigate the overall contribution of C5a signaling to immune cell states, we performed single-cell CITE-seq on FACS-sorted alveolar leukocytes from WT and C5aR1/2^-/-^ mice (n = 4 mice/group) following Spn4 infection. Integrated transcriptomic and surface protein profiling identified major immune cell populations and resolved PMN and AM into 8 distinct subclusters each (PMN 1-8 and AM 1-8; **Fig. 7a**, **Extended Data Fig. 7a**), defined by canonical lineage and activation markers (**Fig. 7b**, **Extended Data Fig. 7b-c**). Less abundant populations included CD8^+^, CD4^+^ and γδ T cells, NK cells, B cells, dendritic cells (DCs), and inflammatory monocytes and DCs (iMoDCs). Consistent with **Fig. 5a-c**, C5aR1/2^-/-^ mice showed increased PMN and relative abundance across multiple subclusters (PMN 1, 2, 4, 6, 7 and 8), alongside reduced AM, most prominently in AM 1 subcluster (∼3-fold) (**Fig. 7c**).

**FIGURE 7.**
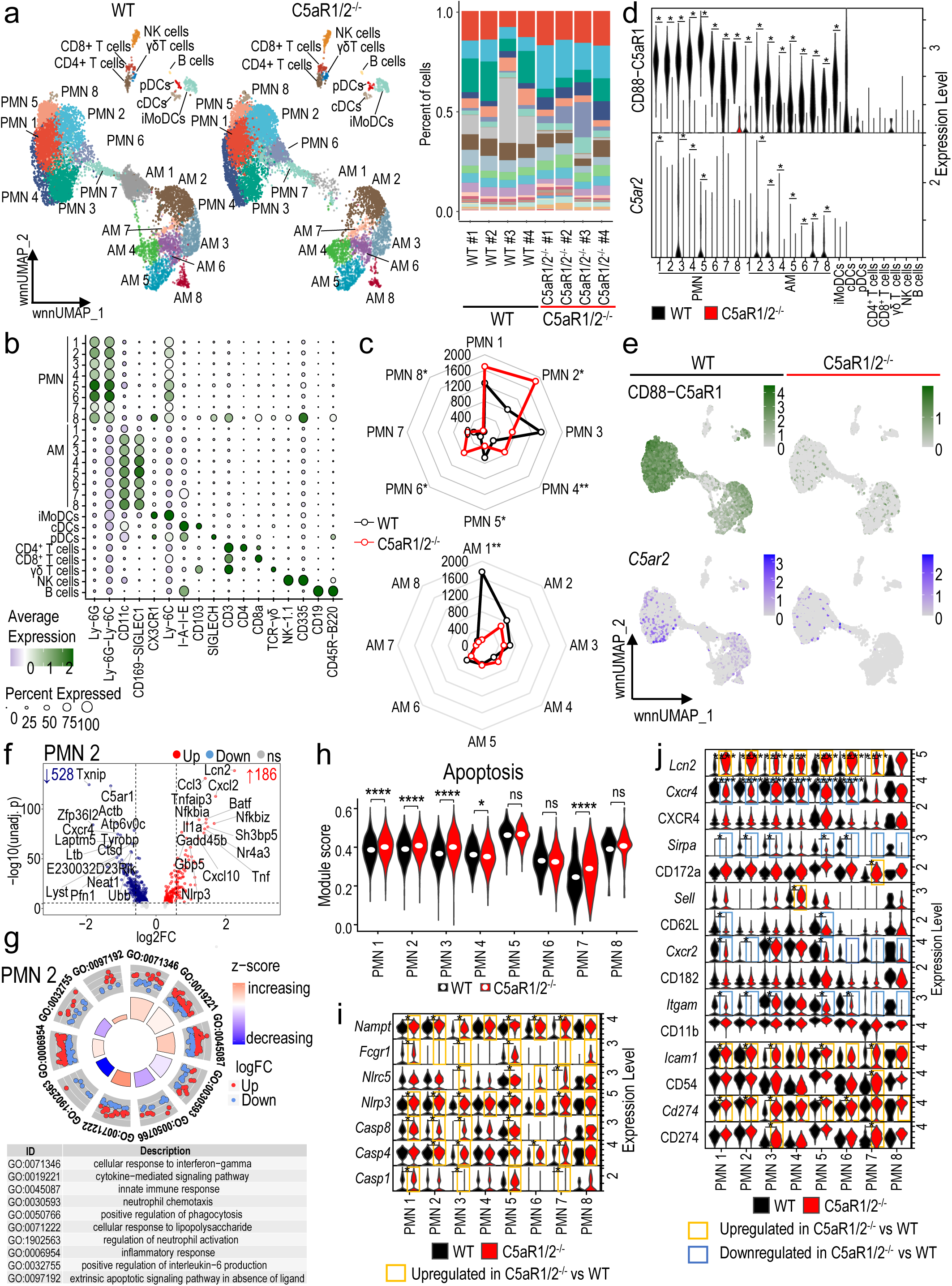
Concomitant deletion of C5aR1 and C5aR2 amplified transcriptomic shifts in alveolar macrophages (AM) and neutrophils (PMN) after *S. pneumoniae*-induced pneumonia in mice. Multimodal sequencing was performed using Cellular Indexing of Transcriptomes and Epitopes by Sequencing (CITE-seq) of FACS-sorted CD45^+^ live BALF cells from WT and C5aR1/2^-/-^ mice (n = 4/group) infected with 2x10^6^ CFU Spn4 for 18 h. (**a**) Plots showing weighted nearest neighbor uniform manifold approximation and projection (wnnUMAP) and cell type distribution in WT and C5aR1/2^-/-^ mice. (**b**) Dot plot showing expression of selected protein markers used for cell type annotation. (**c**) Spider plots showing PMN and AM cluster distributions in more detail. Statistical significance between C5aR1/2^-/-^ vs. WT are indicated next to each subcluster label. (**d-e**) Expression of CD88 (C5aR1) protein and *C5aR2* RNA across cell types visualized using (**d**) violin plot and (**e**) Feature plot. (**f**) Volcano plot of differentially expressed genes in PMN 2 cluster. The number of significant up-/downregulated genes are indicated, with top 15 genes with |log2FoldChange (FC)| > 0.58 labelled. The horizontal dotted line represents adjusted p (adj. p) = 0.05 and the vertical lines represent |log2 fold change| = 1. FC, Fold change; unadj. p, unadjusted p; ns, not significant. (**g**) Circle plot showing selected immune-related gene ontology biological processes (GO BPs) enriched in PMN 2. (**h**) Violin plot showing module score based on apoptosis related GO BPs across PMN clusters. (**i-j**) Violin plot showing expression of key genes functioning in (**i**) apoptosis and cell death, and (**j**) activation and exhaustion in PMN clusters. In panel **j**, italicized names in the y-axis denote RNA expression and non-italicized names protein expression. *adjusted p < 0.05, **p < 0.01, ***p < 0.001, ****p < 0.0001, ns, not significant.

C5a receptor expression was heterogeneous across alveolar immune cells (**Fig. 7d-e**). In WT mice, surface C5aR1 protein was generally higher in PMN than AM, peaking in PMN 5 subcluster. Robust expression was also detected in iMoDCs and cDCs, while expression in pDCs, NK and T cells was minimal. *C5ar2* transcript levels were lower overall but selectively enriched in PMN 1, 3-5 and AM 2-8 subclusters, with little to no expression in other leukocytes. Despite their negligible C5a receptor expression, T and NK cells numbers tended to increase in C5aR1/2^-/-^ mice, suggesting indirect effects that amplify inflammation (**Extended Data Fig. 7d**).

In PMN 2, the most expanded PMN subcluster in C5aR1/2^-/-^ mice, differential expression DEG analysis identified 186 upregulated and 528 downregulated genes (**Fig. 7f**). Upregulated inflammation-associated genes included *Cxcl2*, *Cxcl10*, *Ccl3*, *Tnf*, *Il1a, Nlrp3,* and *Lcn2*, whereas downregulated genes were associated with homeostasis and neutrophil function (*Cxcr4*, *Txnip*), cytoskeletal organization and trafficking (*Actb*, *Pfn1* and *Lyst*), and lysosomal/vesicular processes (*Laptm5*, *Ctsd*, *Atp6v0c*), collectively reflecting impaired effector capacity (**Fig. 7f**). GO analysis revealed enrichment of innate immune, cytokine-mediated, LPS- and IFNγ-response programs, alongside reduced programs associated with regulation of PMN activation, and phagocytosis (**Fig. 7g**). Interferon-associated genes (*Irf1*, *Irf2*, *Irf7* and *Ifnar2*) were also broadly upregulated across PMN subclusters (**Extended Data Fig. 7e**). Extrinsic apoptotic signaling was strongly enriched in PMN2 (**Fig. 7g**), and module scores derived from apoptosis-related GO gene sets suggested increased apoptotic priming across multiple PMN and AM subclusters (**Fig. 7h**, **Extended Data Fig. 7f**). Genes involved in cell death, including inflammatory caspases (*Casp1*, *Casp4*), the apoptotic initiator *Casp8*, inflammasome components (*Nlrp3*, *Nlrc5*), and regulators of activation and metabolic stress (*Fcgr1* and *Nampt*), which have been associated with delayed neutrophil apoptosis, were upregulated across C5aR1/2^-/-^ PMN clusters (**Fig. 7i**). C5aR1/2^-/-^ PMN also exhibited exhaustion-like features - previously described in inflammatory settings^36^, including increased PD-L1/*Cd274* and CD54/*Icam1*, as well as reduced adhesion/migration receptors (CD11b/*Itgam*, CD62L/*Sell*, CD182/*Cxcr2*) at the transcript and protein levels (**Fig. 7j**). Dampened CD172/*Sirpa* suggested altered inhibitory signaling, whereas increased *Lcn2* further indicated dysfunctional inflammatory reprogramming across C5aR1/2^-/-^ PMN subclusters. The global enrichment for inflammatory and apoptotic hallmark pathways extended to myeloid clusters, and highlighted shared IFNγ, TNFα-NFκB, IFNα, IL6-JAK- STAT3 and IL2-STAT5 signaling, but divergent PI3K-AKT-mTOR signaling between AM and PMN populations (**Extended Data Fig. 7g**).

Together, these data indicate that combined C5aR1/C5aR2 deficiency shifts alveolar myeloid populations toward PMN predominance and promotes dysregulated PMN states characterized by hyperinflammation, apoptotic priming, and exhaustion-like features.

### Combined C5aR1/C5aR2 deficiency promotes neutrophil death and exhaustion-like features during pneumococcal pneumonia

To functionally validate the CITE-seq findings, we assessed PMN apoptosis, cell death pathways and exhaustion-associated features during Spn4 infection. C5aR1/2^-/-^ PMN showed increased signals for cleaved Casp-3, a key executioner of apoptosis, (**Fig. 8a-d**). Bax, a pro- apoptotic regulator of the intrinsic apoptotic pathway, was also elevated (**Fig. 8e-h**). Consistently, Annexin V/propidium iodide staining revealed reduced PMN viability, with increased early apoptotic and dead PMN populations in C5aR1/2^-/-^ mice (**Fig. 8i-j**). elevated LDH activity in BALF further indicated increased lytic cell injury within the alveolar compartment (**Fig. 8k**). Despite reduced cell viability, C5aR1/2^-/-^ PMN exhibited increased Ki67 expression, reflected by higher frequencies, absolute counts, and gMFI (**Fig. 8l-o**), which is unlikely to indicate local proliferation but instead may suggest accelerated PMN turnover. In line with this dysregulated state, C5aR1/2^-/-^ PMN displayed exhaustion-like features, including increased PD-L1 expression (**Fig. 8p-s**) and altered surface receptor profiles, with reduced CXCR2 and CD11b alongside increased CD54, indicative of dysregulated migratory and activation capacity (**Fig. 8t-v**). In addition, C5aR1/2^-/-^ AM displayed similar but more modest changes, including increased Bax expression and trends toward elevated cell death and Ki67 expression, with no consistent alterations in PD-L1 (**Extended Data Fig. 8**).

**FIGURE 8.**
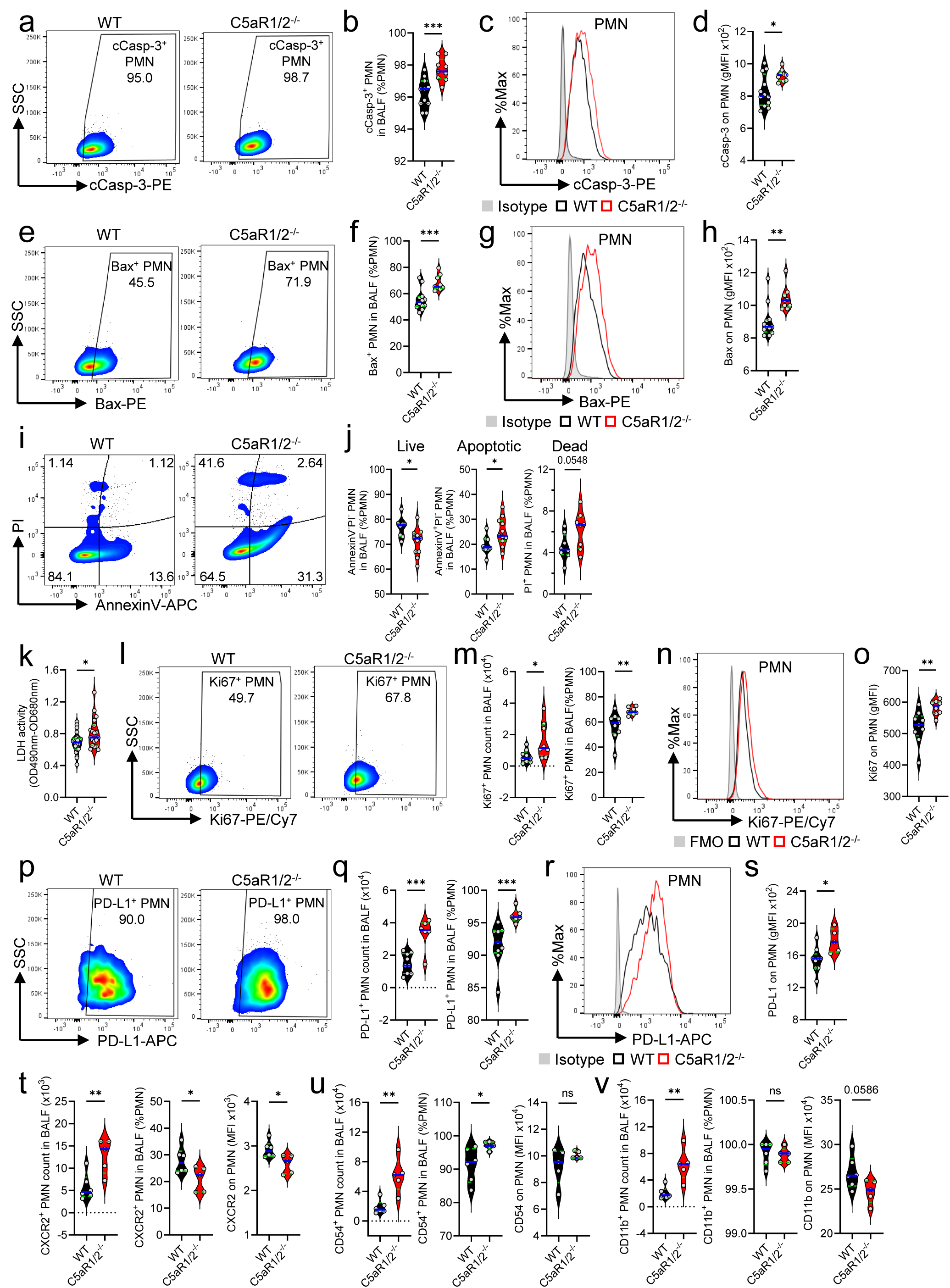
Disruption of C5a-C5aR1/2 axis promotes neutrophil exhaustion and excessive cell death during pneumococcal pneumonia. WT and C5aR1/2^-/-^ mice were intranasally (i.n.) infected with Spn4 (1-2x10^6^ CFU) or PBS (sham), and BALF were collected 18 h later. (**a, e**) Representative flow cytometry plots of cleaved Caspase-3 (cCasp-3) and Bax on Ly6G^+^ neutrophils (PMN) from BAL. Cell frequencies of total PMN are indicated next to each gate. (**b, f**) Percentages of live c-Casp-3^+^ or Bax^+^ total PMN were determined by flow cytometry (n ≥ 9/group). (**c, g**) Representative overlay histograms and (**d, h**) gMFI of cCasp-3 or Bax on PMN. Isotype control was included as a reference (n ≥ 9/group). (**i**) Flow cytometric analysis measuring apoptosis in PMN using AnnexinV and Propidium Iodide (PI) staining. Cell frequencies of total PMN are indicated next to each gate. (**j**) Bar charts show the percentage of live (AnnexinV^−^PI^−^), apoptotic (AnnexinV^+^PI^-^) and dead (PI^+^) PMN in both groups (n ≥ 8/group). (**k**) Pyroptotic cell death was determined by measuring Lactate Dehydrogenase (LDH) in fresh BALF supernatants (n = 19- 22/group). (**l**) Representative flow cytometry plots and (**m**) cell count and fraction of PMN that were actively proliferating, expressed as the percentage of total Ki67^+^ PMN (n = 8-9/group). (**n**) Representative overlay histograms and (**o**) gMFI of Ki67 on PMN. FMO was included as a control (n = 6-9/group). (**p**) Representative flow cytometry dot plots, and (**q**) cell count and fraction of total PMN expressing PD-L1 (n = 5-9/group). (**r**) Representative overlay histograms and (**s**) gMFI of PD-L1 expression on PMN. (**t-v**) Absolute numbers and percentages of total PMN expressing (**t**) CXCR2/CD182, (**u**) CD54/ICAM-1 and (**v**) CD11b determined by flow cytometry. gMFI of listed markers on PMN was also presented. Data are representative of 2-3 independent experiments with similar results. Each circle represents an individual mouse. Horizontal plain blue line across each violin indicates median and green dotted line indicates quartiles. *p < 0.05, **p < 0.01, ***p < 0.001, ****p < 0.0001, as determined by Student’s t-test. BALF, bronchoalveolar lavage fluid; FMO, fluorescence Minus One; gMFI, geometric mean fluorescence intensity.

In summary, these findings support the single-cell proteogenomics profiles and identified a dysregulated maladaptive neutrophil state characterized by increased cell death, sustained recruitment, and functional exhaustion following loss of C5aR1/2 signaling.

## Discussion

Complement activation is a central component of antibacterial host defense, yet the specific contributions of C5a receptors during infection remain incompletely defined. Here, we identify the C5a-C5aR1/2 axis as a regulator of pulmonary host defense during *S. pneumoniae* infection that couples inflammatory response to antimicrobial function in alveolar myeloid cells. Although C5a signaling is widely associated with inflammatory amplification, dual deficiency of C5aR1 and C5aR2 did not attenuate pneumococcus-induced inflammation. Instead, disruption of chemotactic C5a signaling by C5aR1/2 deficiency resulted in excessive PMN accumulation, exuberant NET formation, reduced phagocytic capacity of AM, heightened cytokine production, aggravated lung injury and impaired bacterial clearance. Single-cell profiling linked this phenotype to a dysfunctional PMN state characterized by apoptosis, defective effector capacity, together with features of exhaustion, apoptosis, and broad transcriptional reprogramming across AM and PMN clusters. These findings reveal cooperative but distinct functions of C5aR1 and C5aR2 in balancing inflammation and antibacterial host defense.

C5aR1 has traditionally been considered the principal pro-inflammatory C5a receptor, whereas C5aR2 has been assigned primarily modulatory functions^17, 20, 37^. Our findings support a more nuanced framework in which both receptors exert distinct yet complementary functions during bacterial pneumonia. Single-cell CITE-seq of alveolar immune cells, supported by AM transcriptional profiling, uncovered profound phagocyte dysregulation in infected C5aR1/2^-/-^ mice. Across distinct PMN subclusters, inflammatory and immune effector signatures coexisted with defective antimicrobial programs and features of functional exhaustion. Notably, despite these marked cellular alterations, bulk RNA-seq showed relatively modest transcriptional remodeling in C5aR1/2^-/-^ PMN compared with single KOs, underscoring that heightened inflammatory activation does not necessarily confer effective antimicrobial competence. Complement receptor interplay appears highly context-dependent^38, 39^. During pneumococcal infection, dual loss of both C5aR1 and C5aR2 unmasked cooperative roles in antimicrobial defense and inflammatory regulation, whereas in sterile C5a-induced ALI, C5aR1/2^-/-^ mice phenocopied C5aR1 deficiency, indicating that C5aR1 predominates in the absence of microbial cues. These observations support a context-specific contribution of C5aR2 in host defense, potentially through integrating pathogen-derived signals^13, 40, 41^, modulation of signal intensity, duration or spatial organization^16, 19, 42, 43^. Disruption of this coordination dissociates PMN accumulation from effector function, resulting in impaired bacterial clearance despite excessive neutrophil infiltration, increased tissue injury, and amplified inflammatory programs in AM.

NET formation is a prominent PMN response to infection^44, 45, 46, 47^, but excessive or persistent NET accumulation can promote tissue injury^35, 48, 49, 50^. Accordingly, increased NETosis in C5aR1/2^-/-^ mice was associated with aggravated lung injury, inflammation, and impaired host defense, consistent with previous reports^51, 52, 53, 54^. Increased secreted and extracellular citrullinated histones coincided with reduced suicidal NETosis, and enhanced non-NETotic cell death suggests a shift from canonical lytic pathways toward dysregulated, non-lytic NET release^55, 56^. Both PMN-intrinsic heterogeneity, and extrinsic factors, such as pathogen nature and intercellular signals are likely to determine the preferential engagement of one form of NETosis over the other^47, 48, 52^. NET turnover is also tightly regulated by macrophage-mediated clearance, through which they integrate NET-derived signals to shape antibacterial responses^57, 58, 59^. Notably, suicidal NETosis can potentiate macrophage antibacterial activity against *S. pneumoniae*, rather than directly enhancing PMN-intrinsic microbial killing^60, 61^. Thus, reduced suicidal NETosis combined with impaired macrophage phagocytosis in C5aR1/2^-/-^ mice may create a dual defect in NET-mediated signaling and clearance, sustaining inflammation and tissue injury^58, 59^. Collectively, these findings link dysregulated NET dynamics to persistent macrophage activation and compromised host defense.

Despite robust airway accumulation, C5aR1/2^-/-^ PMN failed to control infection while exhibiting increased apoptosis, transcriptional remodeling, reduced CXCR2 and CD11b (key activation and trafficking receptors), and increased CD54 (adhesion molecule) and PD-L1 (immunosuppressive marker), consistent with exhaustion-like PMN states in severe inflammation^36, 62^. Increased Ki67, elevated BALF GM-CSF and reduced *Cxcr4* expression further suggest that alveolar PMN accumulation may reflect increased turnover and trafficking rather than local proliferation, potentially involving the recruitment of functionally impaired PMN with altered maturation profiles, as reported in sepsis or COVID-19^48, 56, 63, 64^. PD-L1 upregulation was particularly notable, given the association of PD-L1^+^ PMN with immunosuppression and severity of bacterial sepsis^65, 66, 67, 68^. These findings suggest that C5aR1/2 signaling may restrain expansion of PD-L1^+^ PMN, potentially through IFNƴ- dependent regulation^69, 70, 71, 72^. Because PD-L1 expression also correlates with increased NET formation and lung injury^64, 66, 67, 73^, it may link disrupted C5a signaling to PMN dysfunction and impaired bacterial clearance.

The intact alveolar-capillary barrier restricts the passage of large plasma proteins, likely limiting the availability of circulating C5 within the alveolar compartment under homeostatic conditions. Our findings identify AT2 cells as the predominant local source of C5, consistent with our previous transcriptomic analyses showing negligible C5 expression in other pulmonary cell populations^31^. AT2-derived C5 may thus provide a local reservoir for rapid complement activation independently of plasma extravasation. The increased susceptibility of C5-deficient mice to pneumococcal pneumonia with higher neutrophil influx largely phenocopied that of C5aR1/2-deficient mice, underscoring the protective role of C5a signaling in pulmonary host defense.

Our findings underscore the context-dependent role of C5a signaling in pulmonary inflammation. Although C5a promoted lung injury under sterile conditions, disruption of C5a signaling during pneumococcal pneumonia impaired host defense despite increased PMN accumulation. In contrast to previous studies showing protective effects of a C5 blocking antibody or a C5a neutralizing aptamer in pneumococcal infection models^25, 74^, our results caution against considering dual C5a receptor blockade in sepsis and ARDS^74, 75, 76^. Our data further reveal previously unrecognized cooperative and partially overlapping functions of C5aR1 and C5aR2 that only become apparent during pneumococcal infection but cannot be fully appreciated from single-receptor deficiency alone. From a translational perspective, given ongoing international differences in vaccination recommendations for patients receiving C5 inhibitors and pharmacovigilance signals of disproportionately reported pneumococcal pneumonia during eculizumab treatment^77^, our findings provide additional experimental support for broader consideration of pneumococcal vaccination in this population.

## Methods

### Research animals

All studies with mice were approved by the State Investigation Office of Rhineland-Palatinate and the Institutional Animal Care and Use Committee (IACUC) of Boston University. Wildtype (WT) C57BL/6J mice were obtained from The Jackson Laboratory (Bar Harbor, ME). C5aR1^-/-^ mice (B6.129S4-C5ar1tm1Cge) were originally provided by Prof. J. Köhl, Lübeck University. C5aR2^-/-^ (C5ar2tm1b(EUCOMM)Wtsi) mice were generated from targeted embryonic stem cells by our laboratory and the Transgenic Facility of the University of Mainz, Germany. C57BL/6J, C5aR1^-/-^, C5aR2^-/-^, C5aR1/2^-/-^ (all on C57BL/6J background), C5^lacZ^ (B6N(Cg)- Hctm1b(EUCOMM)Wtsi/J) mice and their control C57BL/6N were maintained in a specific pathogen-free environment. All mice were co-housed in a 45-60% humidity, 22 ± 2°C ambient temperature, controlled light/dark (12 h/12 h) cycle, with free access to food and water and were used as sex-age matched cohorts for experiments. Unless indicated otherwise, 8-12- week-old male mice were used for experiments.

### Generation of CRISPR/Cas9 C5aR1/2^-/-^ mice

C5aR1/2 double-knock out (C5aR1/2^-/-^) mice were generated using CRISPR/Cas9 with assistance of the Transgenic Facility of the Johannes Gutenberg University of Mainz, Germany. The commercially available cloning-free CRISPR system from IDT (Integrated DNA Technologies, Coralville, USA) was used to edit the target genes. The CRISPR RNAs (crRNAs) were designed using the CRISPOR tool^78^ as well as the UCSC Genome Browser. The crRNAs with the highest possible score, i.e. low expected off-target effect, were selected.

For mutagenesis, the mastermix for pronucleus/zygote injection was prepared under sterile, RNase-free conditions in RNase-free duplex buffer. Four gRNAs, with two targeting each side of the sequence to be deleted, were combined to increase cleavage efficiency and the likelihood of generating the intended genomic deletion (**Fig. 1a**, **Supplementary Table 1**). The crRNAs were first hybridized individually with a stoichiometric amount of trans-activating CRISPR RNA (tracrRNA) (working concentration 12.5 ng/μL) by heating for 5 min at 95°C and then cooling to room temperature. The resulting gRNA species were pooled at a final concentration of 25 ng/μL. Subsequently, recombinant Cas9 protein was added to the mastermix at a final concentration of 30 ng/μL immediately before microinjection into zygotes. Treated zygotes were cultured for 24 h under sterile conditions before transplantation into sham-pregnant C57BL/6J females. An F0 male founder mouse was identified among >50 offspring mice using PCR-based screening and backcrossed onto C57BL/6J background females for >3 generations to eliminate off-target effects. The newly generated mouse strain C57BL/6J-Del(7C5ar2-C5ar1)1Bosm showed normal lifespan, fertility, organ development, and no other abnormalities in unchallenged mice. The 12.6 kb deletion of the coding regions of C5aR1 (exon 2) and C5aR2 (exon 1 and 2) on chromosome 7 was confirmed by Sanger sequencing.

Genotyping primers are listed in **Supplementary Table 2**. WT band size on PCR1 (R1-P2fo x R1-P1rev): 587 bp and PCR2 (R2-P2fo x R2-P1rev): 913 bp, and mutant band size on PCR3 (R1-P2fo x R2-P1rev): 844 bp (**Fig. 1b**). A faint nonspecific amplification product of ∼350 bp was observed in both genotypes and did not interfere with genotype assignment.

### Whole genome sequencing

To exclude off target mutations in the CRISPR-generated C5aR1/2^-/-^ mouse colony, lung genomic DNA (gDNA) was extracted using the QIAamp Fast DNA Tissue kit (Qiagen), quantified by Nanodrop 2000c, and gDNA quality validated by gel electrophoresis. The whole genome was sequenced on an Illumina Novaseq 6000 platform. Data were analyzed for SNPs and Indels located 20 nucleotides up-/downstream of the gRNA homologous sites as described in **Supplementary Methods**.

### Immunophenotyping of C5aR1/2^-/-^ mice

#### Sample preparation

WT and C5aR1/2^-/-^ male and female mice (n ≥ 5/strain/gender) were weighted weekly from 4 to 16 weeks of age. At 17 weeks old, mice were sedated with isoflurane and blood collected by intracardiac puncture. After euthanasia, a complete necropsy was performed, and organs were isolated, measured and weighted. Spleens were kept in ice-cold PBS until further processing. Blood samples were used for blood cell counts (K-EDTA, Microvette CB300K2E, Sarstedt, Nuembrecht, Germany), clinical chemistry (Lithium-Heparin plasma, Li-He, Microvette 300 LH, Sarstedt) and cholesterol analysis (serum). For blood cell counts, K-EDTA blood was diluted 1:3 in 0.9% (w/v) NaCl. For clinical chemistry, Li-He blood was centrifuged at 2,000 g for 10 min at 4°C and plasma were kept on ice until analysis. For serum preparation, blood was stored for 20 min at room temperature and then centrifuged at 2,000 g for 10 min at 4°C. Sera were further processed for cholesterol determination. The Central Lab of the University of Medicine of Mainz (Germany) performed blood, plasma and serum analysis.

#### Bacterial culture

*S. pneumoniae* serotype 4 strain TIGR4 (Spn4)^79^ were cryopreserved at -80°C using Microbank™ beads (Pro-Lab Diagnostics, Richmond Hill, Canada) as single use aliquots and cultured for intranasal infections as previously described^80^. Briefly, pneumococci were initially grown on 5% blood Columbia agar plates (BBL™, BD Biosciences, Franklin Lakes, USA) overnight in a humidified atmosphere at 37°C in 5% CO_2_. On the day of infection, cultured colonies were transferred to medium containing 3% Todd-Hewitt broth (BD Biosciences) and 0.5% yeast extract (Sigma-Aldrich, St. Louis, USA), and grown to mid-logarithmic phase (optical density of 0.3 at 600 nm wavelength) to obtain ∼1x10^8^ Colony-Forming Units (CFUs) per mL. After centrifugation, the bacterial pellet was resuspended in sterile PBS (Thermofisher, Grand Island, USA) and kept on ice during infection. The infectious dose was confirmed by plating serial dilutions of the inoculum and extrapolation of CFU count after overnight culture.

#### Isolation of splenocytes

Single-cell suspensions from spleens were prepared using an adapted protocol described by Stagg et al.^81^. Briefly, spleens were minced after organ injection of 1 mL of digestion medium containing RPMI-1640, 10% heat-inactivated FBS (hi-FBS), 0.02 mg/mL DNase I (AppliChem, Darmstadt, Germany) and 1 mg/mL Collagenase D (Roche Diagnostics, Mannheim, Germany), and gently agitated for 1 h at 37°C. The digestion was stopped by adding EDTA to reach 10 mM final concentration. The digested samples were filtered through a 100 μm cell strainer and centrifuged at 1350 rpm for 5 min at 4°C. Red blood cells were lysed by resuspension in 1 mL Ammonium-Chloride-Potassium (ACK) lysis buffer (Life Technologies) for 3-4 min at room temperature. Next, 9 mL cold complete medium (RPMI -1640 supplemented with 10% hi-FBS) were added to stop lysis followed by centrifugation at 1,350 rpm for 5 min. Cells were washed twice and resuspended in 5 mL complete medium and kept on ice until flow cytometry staining. For leukocyte profiling, 100 μL of the splenocyte suspension was used per antibody panel (**Supplementary Table 3**). The gating strategy presented in **Extended Data Fig. 2** was established according to current flow cytometry guidelines^82^.

#### Lung injury and bacterial pneumonia

Mice were anesthetized by i.p. injection of Ketamine (100 mg/kg body weight) and Xylazine (8 mg/kg body weight).

Acute lung injury (ALI) was induced by intratracheal instillation of rmC5a as described before^30,35^. Briefly, the cervical region was disinfected, incised and trachea was carefully dissected with blunt forceps. Recombinant mouse C5a (rmC5a, 500 ng/mouse, R&D Systems, Minneapolis, USA) resuspended in 40 μL of sterile PBS was slowly instilled into the trachea using a microliter syringe (Hamilton company, Reno, USA) mounted with a 30G needle, while control animals were instilled intratracheally with PBS only. Mice were maintained in a vertical position for one minute to allow proper inoculum distribution before wound closure with surgical clips.

For bacterial pneumonia, mice were challenged with 1-2x10^6^ pneumococci in 20 μL of PBS by intranasal (i.n.) application. The body weight was monitored daily and ≥ 20% weight loss or severe distress were defined as humane endpoints for euthanasia.

At the end of experiments, mice were euthanized by CO_2_ asphyxiation, and BALF were collected in 1 mL ice-cold sterile PBS supplemented with 2 mM EDTA and 1X proteases inhibitor (EASYpack Complete Mini EDTA-free cocktail tablet, Roche, Wilmington, USA) by inserting an 18G angiocatheter (Exel International, Redondo Beach, USA) into the trachea.

Bacterial loads were assessed by plating BAL on blood agar plates overnight and counting colony-forming units (CFUs). The lavages were centrifuged at 650 g for 10 min and cell free- supernatants (650 g, 10 min) were stored at -80°C. BALF cell pellets were subjected to red blood cell lysis by resuspension in 200 μL ammonium-chloride-potassium lysis buffer (Life Technologies) for 3 min on ice. Next, 800 μL ice-cold PBS was added to stop lysis followed by centrifugation at 650 g for 10 min.

#### Phagocytosis assay

To assess the ability of alveolar macrophages (AM) to bind Spn4, we used a flow cytometry phagocytosis assay adapted from a previous protocol^83^. In brief, Spn4 were fluorescently membrane-labelled using CellVue Claret dye (Sigma) for 3 min, washed thoroughly in 1% BSA PBS and resuspended in sterile PBS. Mice were challenged i.n. with the fluorescently labeled Spn4 for 90 min before BALF samples were collected and processed for flow cytometry as described above.

#### Quantification of proteins

Total protein amount in cell-free BALF was quantified using Pierce™ BCA Protein assay kit (Thermo Scientific, Rockford, USA). Optical densities were measured with an Infinite M Nano (Tecan, Maennedorf, Switzerland) microplate reader at 562 nm corrected with the absorbance values of the blank reference.

Citrullinated Histone 3 (citH3), a biomarker of NET formation, was measured using Cell Death Detection ELISA^plus^ kit (Roche) following an adapted protocol previously described^53^. Briefly, an ELISA plate was coated with a rabbit polyclonal anti-CitH3 antibody (citrulline R2+R8+R17, 4 μg/mL, Abcam) diluted in carbonate-bicarbonate buffer. 20 μL of cell-free BALF were added to 80 μL of immunoreagent containing the detection anti-DNA antibody (supplied in the kit) but not the capture antibody. Next steps were completed as per the manufacturer’s protocol. Absorbances at 450 nm were measured with an Infinite M Nano (Tecan).

Mouse CXCL10/IP-10, IL-6, TNF-α, IL1-β, CXCL5, CXCL1/KC, MIP-1α/CCL3 and MCP-1/CCL2 ELISA kits were purchased from R&D Systems. Mouse IFN-γ and Albumin ELISA kits were purchased from BioLegend and Bethyl Laboratory (Montgomery, USA), respectively. Cell-free BALF supernatants were analyzed according to the manufacturers’ protocol. In short, samples were diluted in 0.1% BSA in PBS to fit into the standard range. Optical densities of oxidized TMB (Invitrogen, San Diego, USA) were measured with an Opsys MR Dynex (Aspect Scientific, Little Budworth, UK) or Infinite M Nano (Tecan) microplate reader at 450 nm corrected with the reference wavelength of 570 nm.

Cell death was assessed in fresh cell-free BALF supernatants using CyQUANT™ LDH Cytotoxicity Assay (Thermofisher, #C20301).

#### Multiplex bead-based immunoassay

A multiplex bead-based immunoassay (Cytokine & Chemokine 26-Plex Mouse ProcartaPlex™, Invitrogen) was used for simultaneous quantification of 26 cytokines/chemokines from cell-free BALF samples. Eotaxin (CCL11), GM-CSF (CSF-2), GROα (CXCL1), IFN-γ, IL-1β, IL-2, IL-10, IL-12p70, IL-13, IL-17A (CTLA-8), IL-18, IL-22, IL-23, IL-27, IL-4, IL-5, IL-6, IL-9, IP-10 (CXCL10), MCP-1 (CCL2), MCP-3 (CCL7), MIP-1α (CCL3), MIP-1β (CCL4), MIP-2α (CXCL2), RANTES (CCL5) and TNF-α concentrations were measured according to the manufacturer’s instructions. BU Analytical Instrumentation Core performed the Luminex/Drop Array assay. Fluorescence intensities were recorded on a Luminex^®^ Magpix reader (Thermo Fisher Scientific) and analyzed using xPONENT^®^ software (Invitrogen).

#### Flow cytometry

Following red blood cells lysis, BAL cells were washed with ice-cold PBS (650 g, 5 min, 4°C) and dead cells were labeled for 20 min at 4°C with eFluor 780 Fixable Viability Dye (eF780- FVD) using heat-killed (5 min, 65°C) cells as positive controls. After two washes with FACS buffer (1% (w/v) BSA (Fisher Bioreagents, Fair lawn, USA), 0.01% (w/v) sodium azide (Sigma), 2.5 mM EDTA in sterile PBS), cells were preincubated for 20 min at 4°C with anti-mouse CD16/CD32 antibody (TruStain FcX) in FACS buffer to block Fc receptors followed by 20 min incubation at 4°C with appropriate surface fluorescence dye-conjugated anti-mouse antibody cocktail or their isotype controls (**Supplementary Table 3**). Cells were washed twice with FACS buffer, fixed in 2% PFA (Santa Cruz Biotechnology, Dallas, USA) for 20 min at 4°C, washed again twice and resuspended in FACS buffer before acquisition.

For intracellular detection of C5aR2 mRNA expression, the whole lungs were digested as described in the next section. Single-cell suspensions were stained using the PrimeFlow RNA Assay Kit (Thermo Fisher Scientific, Waltham, USA) including β-Actin mRNA as a positive control.

For apoptosis assay, cells were stained with surface antibodies then washed as described above and resuspended in Annexin V Binding Buffer (BD Pharmingen) containing APC- conjugated Annexin V. After 15 min incubation at room temperature, additional Annexin V Binding Buffer was added to the cells as well as Propidium Iodide, and cells were quickly analyzed via flow cytometry within 5 min. For intracellular staining of proliferation marker Ki67 or the apoptosis markers, Bax and cleaved Caspase-3 (cCasp-3), BALF cell suspensions were prepared as described above. After surface staining, cells were fixed and permeabilized using the eBioscience FoxP3/TF staining buffer set. Cells were further incubated with unconjugated anti-Bax or anti-cCasp-3 or PE-Cy7-labelled anti-Ki67 antibodies (or appropriate isotype control) for 30 min at 4°C. Cells were washed twice, stained for 30 min at 4°C with secondary anti-rabbit PE-coupled antibody, washed again twice, resuspended in FACS buffer, and analyzed by flow cytometry.

For citH3 detection, BALF cells were stained with Zombie UV FVD in PBS for 20 min and fixed in 2% PFA prior to cell surface staining including a cocktail of fluorochrome-labeled antibodies and unlabeled primary anti-citH3 antibody or its isotype control. After 20 min incubation at 4°C, cells were gently washed once, incubated with a secondary AF488 anti-rabbit IgG for 20 min at 4°C, resuspended in FACS buffer, and analyzed by flow cytometry.

For cell counting, 123count eBeads (Thermo Fisher Scientific, 1:5) or CountBright™ Absolute counting beads (Invitrogen, 1:15) were added according to the manufacturer’s instructions. At least 5,000 beads or events of interest were acquired with a BD FACSCanto™ II (BD Biosciences) or BD LSR II SORP (BD Biosciences) using BD FACSDiva software. Fcs files were analyzed using FlowJo v10 Software (FlowJo, Ashland, OR).

#### Flow cytometric LacZ/β-galactosidase reporter assay

Lungs were perfused *in situ* with 10 mL PBS via the right heart ventricle after a small incision of the left ventricle. A 18G IV catheter (Introcan-W, B. Braun, Melsungen, Germany) was inserted into the trachea and secured with a suture for sequential infusion of Dispase II (37°C, 45 U/mL, Sigma) and 1% low-melting-point agarose (SeaPrep, Lonza, Rockland, USA), followed by cooling to solidify the agarose. Next, lungs were collected and submerged into prewarmed Dispase II for 45 min at room temperature with gentle shaking. The digested lungs were decanted into a 10 cm Petri Dish and minced in complete medium containing DMEM (Life Technologies), 10% hi-FBS, 100 U/mL Penicillin-Streptomycin, supplemented with 3.5 μg/mL DNase I using forceps. After 10 min of gentle shaking at room temperature, the cell suspension was filtered through 100 μm and 30 μm cell strainers, and centrifuged at 500 g for 10 min at 4°C. RBC were lysed twice in ACK lysis buffer for 3 min on ice and lung cells were washed twice in complete DMEM. Following counting in a Neubauer hemocytometer, 2 x 10^5^ cells per condition were stained with eF780-FVD and antibody panels (**Supplementary Table 3**). To detect LacZ-associated β-galactosidase activity, cells were incubated with the cell-permeable fluorogenic substrate DDAOG (10 μg/mL; Thermo Fisher Scientific) for 1 h at 37°C. DDAOG is cleaved by β-galactosidase to the far-red fluorescent product DDAO (Ex/Em ∼645/660 nm), which is readily detected by flow cytometry with minimal interference from uncleaved DDAOG (Ex/Em ∼ 460/610 nm) or cellular autofluorescence^32^. Finally, cells were washed twice in FACS buffer, fixed in 2% PFA, washed twice again before flow cytometric acquisition.

#### Immunohistochemistry

For paraffin embedding, lungs were transcardially perfused with PBS and airways inflated with 50% ROTI^®^ Histofix (Carl Roth, Karlsruhe, Germany) in PBS, and fixed (50% ROTI^®^ Histofix supplemented with 0.25% Glutaraldehyde and 0.01% NP-40) for 2 h at 4°C. Next, lungs were washed for 3 h at 4°C and stained using 1 mg/mL X-Gal (Boehringer, Mannheim, Germany) containing 5 mM K3Fe, 5 mM K4Fe, 2 mM MgCl2 and 0.02% NP-40 for 24 h at 37°C. Subsequently, lungs were washed in cold PBS for 30 min and fixed again overnight at 4°C in 100% ROTI^®^ Histofix. Lungs were then dehydrated with graded isopropanol, cleared in xylene using a Leica TP1020 automated tissue processor (Leica Biosystems, Danvers, USA), and paraffin-embedded using a Leica EG1150 C&H embedding system. Paraffin blocks were processed for IHC at the Histology Core Facility, University Medical Center Mainz. Sections (2-5 µm) were stained with an anti-mouse CD45 primary antibody, followed by Diaminobenzidine (DAB) detection and Nuclear Fast Red counterstaining (Sigma). Kidneys from wild-type mice were used as positive controls, and sections processed without primary antibody as negative controls. Bright-field images were acquired using Olympus or Zeiss microscopes with 20x and 40x objectives.

For cryo-embedding, lungs were inflated with 50% Tissue-Tek^®^ OCT™ medium (Sakura Finetek, Alphen aan den Rijn, The Netherlands) before collection and fixation in 50% ROTI^®^ Histofix for 1-2 h at 4°C. Other organs were fixed in 100% ROTI^®^ Histofix for 24 h. Tissues were thoroughly washed in cold PBS and cryoprotected sequentially in 15% and 30% sucrose/PBS for ≥ 24 h each at 4°C. Tissues were immersed in OCT for 1 h at 4°C, transferred to OCT-filled molds, and snap-frozen in liquid nitrogen vapor. Frontal sections of lungs and longitudinal or transverse sections of other organs (5-10 µm) were prepared by cryostat (Leica, Wetzlar, Germany) and stored at -80°C until staining. Sections were washed 3 times for 5 min in cold PBS and stained with 1 mg/mL X-Gal (Boehringer) containing 5 mM K3Fe, 5 mM K4Fe and 2 mM MgCl2 for 24 h at 37°C. Slides were washed twice for 5 min in cold PBS, counterstained with Nuclear Fast Red or H&E, dehydrated through graded alcohols, cleared in xylene substitute, and coverslipped with Eukitt® mounting medium (Sigma) before overnight drying. Bright-field images were captured using Olympus or Zeiss microscopes with 5x, 10x or 20x objectives.

#### Confocal immunofluorescence histology

The frozen sections were dehydrated and washed as described above followed by staining with 200 µL of pre-warmed 10 μM DDAOG (containing 25 mM Bicin, 0.25% Triton X-100, 20 mM NaCl and 1 mM MgCl2 in distilled water) for 24 h in the dark at 37°C in a humid chamber. Sections were washed twice for 5 min in PBS supplemented with 0.05% Tween 20, incubated in blocking buffer (2% BSA, 0.05% Tween 20 in PBS) for 30 min at room temperature, and incubated with unconjugated anti-mouse SP-C antibody (rabbit monoclonal IgG, clone EPR19839, 1:100, Abcam, #ab211326) diluted in 0.05% Tween 20 in PBS overnight at 4°C. The sections were washed 3 times for 5 min in 0.05% Tween 20 in PBS, incubated with secondary goat anti-rabbit IgG AF488 antibody (Invitrogen, 1:100, A11008) for 30 min at room temperature, and washed 3 times for 5 minutes in 0.05% Tween 20 in PBS. Sections were stained for nuclei with 200 µl of DAPI (1:20,000) for 3-10 min, washed twice for 5 minutes in cold water, coverslipped with an aqueous mounting medium and allowed to harden overnight in the dark at room temperature. Secondary antibody only-stained sections served as negative controls. All the images were acquired on a Leica TCS SP8 with HyD microscope (20x objective) at a resolution of 1,024 x 1,024 pixels.

#### Ex vivo imaging of LacZ/β-galactosidase activity

Mice were anesthetized, injected i.v. with 200 μL of 2 mM DDAOG or vehicle (DMSO/PBS) and euthanized within 30 minutes. Organs were collected and imaged *ex vivo* using an IVIS Spectrum system (PerkinElmer, Shelton, USA) with DDAO (Ex/Em ∼640/700 nm) and DDAOG (Ex/Em ∼465/600 nm) filter settings. Regions of interest were drawn around each organ, and total radiant efficiency was quantified in both channels.

#### Reverse transcription and real-time qPCR

Total RNA was extracted from isolated cells or tissue (20 mg) with the RNeasy Mini Kit (Qiagen) or innuPREP RNA Mini Kit (Analytik Jena). RNA was quantified using a NanoDrop™ 2000c spectrophotometer (Thermo Fisher Scientific) and reverse-transcribed into cDNA with the High-Capacity cDNA Reverse Transcription kit (Applied Biosystems, Waltham, USA) or the Maxima™ H Minus cDNA Synthesis Master Mix kit (Thermo Fisher Scientific).

For real-time quantitative PCR, 1-2 ng cDNA template was complemented with iQ SYBR® Green Mastermix (Bio-Rad) or PowerUp™ SYBR™ Green Master Mix (Applied Biosystems, Waltham, USA) and primer pairs (500 nM each; **Supplementary Table 4**). Reactions were run on a C1000 Thermal Cycler (BioRad) or a QuantStudio 3 Real Time PCR System (Applied Biosystems), and expression was normalized to *β-actin* using the 2^-ΔΔCt^ method.

#### Bulk RNA-seq library preparation and data analysis

BALF cells were prepared following the same protocol as for flow cytometry. Alveolar macrophages (AM) were defined as FVD^-^CD64^+^Ly6G^-^SiglecF^+^CD11c^+^ singlets and FVD^-^ CD64^-^Ly6G^+^SiglecF^-^CD11c^-^ singlets as neutrophils (PMN) during FACSorting (BD FACSAria™ II SORP, BD Biosciences). Total RNA was extracted using the Qiagen RNeasy Mini Kit (QIAGEN, Germantown, USA) or InnuPREP RNA Mini kit (Analytik Jena, Berlin, Germany). RNA abundance was quantified using a Qubit RNA HS Assay kit (Life Technologies, Thermo Fisher Scientific) on a Qubit fluorometer (Life Technologies). RNA integrity was evaluated using RNA 6000 Nano or Pico total RNA kits (Agilent, Santa Clara, USA) on an Agilent 2100 Bioanalyzer (Agilent Technologies). All samples showed a RIN value of ≥ 8. The RNA-seq libraries were constructed with NEBNext Low-Input RNA Library Prep Kit for Illumina with an extra SMARTer RNA amplification step for neutrophils and sequenced with paired-end reads on a PE150 Illumina NovaSeq 6000 (Novogene, Yuen Long, Hong Kong). The bulk RNA-seq data analysis is described in Supplementary Methods.

### Single-cell RNA-seq (CITE-seq/TotalSeq)

#### Sample preparation

Following Fc receptor blocking (TruStain FcX™ PLUS, BioLegend) in 1X MACS buffer (2.5 mM EDTA, 1% BSA, PBS) for 10 min at 4°C, cells were stained with eF780-FVD and anti- CD45 antibody for 20 min at 4°C. Singlets that were FVD^-^CD45^+^ were sorted using an BD FACSAria™ II SORP (BD Biosciences), and incubated with an anti-mouse TotalSeq™-A pretitrated cocktail of 198 antibodies (custom panel, BioLegend, **Supplementary Table 5**) for 30 min at 4°C according to manufacturer’s instructions. An additional TotalSeq™-A ‘Spike-in’ antibody (TotalSeq™-A1042 anti-mouse CD88, clone 20/70, BioLegend) was added to the staining panel following an optimized protocol for C5aR1 protein. Next, cells were washed 3 times in 1X CITE-seq buffer (0.04% BSA in PBS) and centrifuged (650 g for 10 min at 4°C). Cell viability was assessed for each sample using Trypan Blue (1:1) and a Neubauer hematocytometer (Hausser Scientific), and the few samples that fell below 90% viability were not processed further.

#### 10X processing and library preparation

A target of ∼3,000 per sample was captured using a Chromium Next GEM single-cell 3’ Gel Bead kit v3.1 on each well of a Chip G (10X Genomics). Antibody-Derived Tag (ADT)- and mRNA-derived cDNA libraries preparation, from reverse transcription to final amplification, was performed with a KAPA HiFi Hotstart ReadyMix PCR kit (KAPA Biosystems) and a Chromium Next GEM single-cell 3′ GEM library kit v3.1 (10X Genomics) respectively, following manufacturers’ instructions. For ADT libraries, additional primers were included at cDNA amplification step (ADT Additive Primer) and library construction steps (SI PCR primer and Small RNA RPI1 to RPI8 primers, index i7) as recommended by BioLegend. Quality control of generated intermediate libraries was performed using a High Sensitivity DNA chip run on a Bioanalyzer 2100 device (Agilent). ADT libraries contained a predominant single peak at 180 bp whereas mRNA-derived cDNA libraries were featured by a single peak at ∼500 bp.

mRNA-derived cDNA libraries were indexed using a Single Index kit (Single Index T set A,10X Genomics) and further combined with ADT libraries at desired proportions (10% and 90% of a lane, respectively) for each sample. Pooled Gene Expression (mRNA-derived cDNA) and Cell Surface Protein (ADT) libraries were paired-end sequenced on a NextSeq2000 device (Illumina) at a depth of 1.33 billion reads (92% > 30 and 0.12% error rate), resulting in a read depth of 55,000 reads per cell on average (5,000 reads per cells for ADTs and 50,000 reads per cell for mRNA-derived cDNAs).

#### Raw data analysis

The CITE-Seq/TotalSeq data for the 8 samples (WT and C5aR1/2^-/-^; n = 4 mice/group) were analyzed following standard protocol and are described in the Supplementary Methods.

### Statistical analysis and data presentation

All statistical analyses and graphics were made using GraphPad Prism 10 software or R (The R Project for Statistical Computing). The sample sizes of *in vivo* experiments are provided in the figure legends, and several independent experiments were pooled when possible. Unless otherwise stated, all *in vivo* and *in vitro* experiments were repeated at least three times (biological replicates). ELISA and qPCR were run in duplicate wells (technical replicates). Values are depicted as mean with error bars representing SEM and significance was analyzed using the unpaired two-tailed Student t-test or one-way ANOVA if more than 2 groups were compared. Survival analysis p-values were calculated using the log-rank Mantel-Cox test. Normality and equal variances were formally tested to ensure the data met the assumptions of the statistical tests used. P-values ≤ 0.05 were considered statistically significant in all comparisons. *p ≤ 0.05, **p ≤ 0.01, ***p ≤ 0.001, ****p ≤ 0.0001, ns: not significant (p > 0.05).

## Data availability

All bulk RNA-seq and CITE-seq/TOTALSeq data have been deposited in the Gene Expression Omnibus under the accession numbers GSE297359 (C5a-ALI; reviewer access token: ydstmwmkzxoflqx), GSE297357 (AM Spn4: C5aR1^-/-^, C5aR2^-/-^, C5aR1/2^-/-^; apotiaoulngddsx), GSE297362 (PMN Spn4: C5aR1^-/-^, C5aR2^-/-^, C5aR1/2^-/-^; wxetwoeiffsfnop), and GSE297356 (CITEseq: obexsoymrpqfjyn). In addition, CITE-seq/TOTALSeq data can be further explored under https://shiny.imbei.uni-mainz.de/iSEE_complement/ using the interactive SummarizedExperiment Explorer (iSEE) interface^84^. The whole genome sequencing data generated in this study from C5aR1/2^-/-^ mouse lungs are available in the NCBI SRA repository under Bioproject accession number PRJNA1142148. Further inquiries can be directed to the corresponding author.

## Acknowledgments

We thank Leonid Eshkind from the Transgenic Facility of University of Mainz for his significant contribution to the generation of C5aR1/2^-/-^ mice. We thank Foruzandeh Samangan and Melissa Pesta for their technical assistance, and the members of the Mizgerd laboratory for helpful discussions, including Dr. Anukul Shenoy for his technical contribution to the Spn labelling protocol. We cordially thank Kevin Friedmann and Prof. Sven Danckwardt for help with the IVIS studies, and Dr. Amrit Mann, Dr. Sebastian Rosigkeit and Dr. Ernesto Bockamp for technical expertise in histology and microscopy. C.R. is a member at the German Center for Cardiovascular Research (DZHK), the DFG Research Unit 5644 INFINITE (RE 3450/15-1), the Center for Translational Vascular Biology, the Research Center for Immunotherapy (FZI), and Potentialbereich EXPOHEALTH and received a Fellowship from the Gutenberg Research College. We extend our acknowledgments to the Flow Cytometry cores of the University of Medicine of Mainz and Boston University, especially Dr. Anna Belkina, Brian Tilton and Shari Brezinsky, and the Single Cell Sequencing Core of Boston University. We thank Dr. Lucien Garo and Dr. Filiz Korkmaz for reading the manuscript. The authors are responsible for the contents of this publication. Some data from this paper were used by V.T. for her Doctor of Medicine thesis. All requests for materials and correspondence should be addressed to M.B.

## Author contributions

S.W. and V.T. designed and performed experiments, S.W., V.T., A.J., N.C. and F.M. analyzed data, S.X. managed mice genotyping, S.W. and A.J. wrote the manuscript, M.D. provided protocols and data, M.B. conceived and supervised the study, designed experiments and interpreted data, and provided funding.

## Supplementary information

**Supplementary Tables 1-5 and a supplementary file containing an extended description of the whole genome sequencing, bulk RNA-seq and CITE-seq methods are provided according to the journal’s guidelines.**

## Extended data legends

**Extended Data Fig. 1: Breeding statistics and blood analysis of C5aR1/2^-/-^ double knockout mice.**

Related to Fig. 1. (**a-c**) Analysis of breeding data collected over a 24-month period from inbred mice of C5aR1/2^-/-^ and C57BL/6J wild type (WT) strains. (**a**) Frequency of genders and genotypes of mice born in each configuration of breeding cages. The upper panel depicts the sex ratio of pups born from homozygous (Homo x Homo) breeders, the two charts below show the genders and genotypes distribution across pups born from the breeding of heterozygous and homozygous (Het x Homo) mice, and the lowest two panels represent similar distribution in heterozygous breeding pairs (Het x Het). The number of litters and offspring corresponding to each breeding condition is indicated in the legend of each chart. Unknown genotype frequencies represent offspring mice that were not sampled for genotyping. (**b**-**c**) Analysis of reproductive performance metrics of pair-bred (duo) and trio-bred (trio) mice for each category of breeding condition. (**b**) Bar graphs represent the absolute frequencies (number of Iitters) across litter size categories. The median of litter size is presented in blue and the fit curves show the frequency distribution of litter size for each breeding configuration. The corresponding total number of litters is stated below each graph. The right panel summarizes the average litter size in each duo-bred cage. (**c**) Number of days between births in each breeding configuration. Values are represented as mean ± SEM. (**d**-**k**) C5aR1/2^-/-^ mice (females: ♀; males: ♂) and their C57BL/6J (WT) counterparts were (**d**) weighted weekly from 4-16 weeks after birth. (**e**) Mice were necropsied at 17 weeks of age and organ weights were recorded. (**f-h**) Clinical chemistry analysis of peripheral blood. (**i**) Quantification of lipids. (**j-k**) Differential blood cell count. Bar graphs show mean ± SEM values from the analysis of n = 5 mice per strain and per gender. Data are representative of 3 independent experiments. *p < 0.05 as determined by Student’s t-test analysis. GPT, Alanine aminotransferase; GOT, Aspartate aminotransferase; LDH, Lactate Dehydrogenase; CRP, C Protein Reactive; HDL, High Density Lipoprotein; LDL, Low Density Lipoprotein; LUC, Large unstained cells.

**Extended Data Fig. 2: Immunophenotyping analysis of splenocytes from C5aR1/2^-/-^ double knockout mice.**

Related to Fig. 1. C5aR1/2^-/-^ mice (females: ♀; males: ♂) and their C57BL/6J (WT) counterparts were necropsied at 17 weeks of age and spleens were collected to isolate immune cells by enzymatic digestion and analyzed by flow cytometry. (**a, c, e, g**) Representative gating strategy for splenic immune cells in naïve mice. After doublets and dead cells exclusion (eF780-FVD^+^ cells), further gating was used to identify, (**a**) Lymphocytes (T cells, CD8^+^ and CD4^+^ T cells, B cells, NK cells, ILC1 cells), (**c**) Leukocytes (CD45^+^ cells), Granulocytes (Neutrophils, Eosinophils, Basophils), (**e**) Monocytes/Macrophages, and (**g**) Dendritic cells (pDC, cDC, cDC1, cDC2). Lineage: CD3/CD19/CD49b/Ly6G. (**b, d, f, h**) Graphs show total immune cell counts in each spleen. Bars show mean ± SEM values from the analysis of n ≥ 5 mice per strain and per gender. Data are representative of 3 independent experiments. *p < 0.05 as determined by Student’s t-test analysis. FVD, Fixable Viability Dye; NK, Natural Killer; ILC1, Innate Lymphoid cells 1; pDC, plasmacytoid DC; cDC, conventional DC; lo, low; int, intermediate; hi, high.

**Extended Data Fig. 3: Transcriptional regulation of inflammatory mediators and genes involved in tissue remodeling and healing processes in alveolar macrophages (AM) following C5a-ALI in WT mice.**

Related to Fig. 1 and 2. (**a**) Representative gating strategy, (**b**) cell frequencies and (**c**) absolute cell counts of BALF infiltrating immune cell populations 7 h after rmC5a (500 ng) or PBS (sham) i.t. instillation in WT, C5aR1^-/-^, C5aR2^-/-^ and C5aR1/2^-/-^ mice identifying Ly6G^+^Ly6C^+^ neutrophils (PMN), SiglecF^+^CD11c^+^ alveolar macrophages (AM), Ly6G^-^Ly6C^+^ monocytes (Mono) and SiglecF^+^CD11c^-^ eosinophils (Eos). Inset numbers indicate percentages of cells from parental gates falling into daughter gates. (**b-c**) Data are shown as mean ± SEM and are representative of 2-3 independent experiments. Each circle represents an individual mouse; (**b**) n = 6-15 mice/group, (**c**) n = 4-11 mice/group. *p < 0.05, **p < 0.01, ***p < 0.001, ****p < 0.0001, ns, not significant, as determined by one-way ANOVA. rmC5a, recombinant mouse C5a; BALF, bronchoalveolar lavage fluid. (**d-e**) Bulk RNA-seq was performed on FACS- sorted live SiglecF^+^CD11c^+^CD64^+^ AM from sham and C5a-ALI mice (n = 6-8 mice/group). Heatmaps showing significant DEGs enriched in gene ontology biological processes (GO BPs) associated with (**d**) tissue remodeling and wound healing; (**e**) production, regulation, and response to inflammatory mediators (cytokines and chemokines) and reactive oxygen species metabolism. Heatmaps visualize normalized expression values of DEGs with baseMean ≥ 1000 with Z-score from -2/blue to +2/red. SigF, SiglecF.

**Extended Data Fig. 4: Identification of C5-producing cells in lungs of naïve reporter mice.**

Related to Fig. 3. (**a-b**) Representative images of immunohistochemistry analysis for X-Gal and Nuclear Fast Red (**a**, left and middle panels) or H&E (**a**, right panels and **b**) staining of fixed-frozen liver (**a**) and indicated organs (**b**) sections from C5^lacZ^ and WT control mice. (**a**, left and middle panels) Scale bars = 5 mm (whole section), 200 μm (zoomed sections). (**a**, right panel) Scale bars = 500 μm. (**b**) Scale bars are indicated under the respective panels. The data are representative of 3 experiments with similar results. (**c**) Dot plots showing the representative gating strategy used for alveolar type 2 epithelial cells (AT2) in enzymatically digested lungs of naïve C5^lacZ^ mice by flow cytometry. Cell debris, doublets and dead cells were excluded prior to lung single-cell suspensions analysis. Pdpn, podoplanin.

**Extended Data Fig. 5: Loss of C5aR1 and C5aR2 differentially remodels inflammatory pathways in alveolar macrophages during *S. pneumoniae* infection in mice.**

Related to Fig. 6. Bulk RNA-seq was performed on FACS-sorted live SiglecF^+^CD11c^+^CD64^+^ alveolar macrophages (AM) from WT, C5aR1^-/-^, C5aR2^-/-^ and C5aR1/2^-/-^ mice 18 h after intranasal infection with 2x10^6^ CFU Spn4. Gene set enrichment analysis (GSEA) showing selected immune-related hallmark pathways significantly altered in C5aRs-deficient AM (adjusted p < 0.05) based on the analysis of significant and non-significant differentially expressed genes (DEGs). NES, normalized enrichment score.

**Extended Data Fig. 6: Single or combined deletion of C5aR1 and C5aR2 impaired neutrophil immune responses during pneumococcal infection in mice.**

Related to Fig. 6. Bulk RNA-seq was performed on FACS-sorted live Ly6G^+^ alveolar neutrophils (PMN) from WT, C5aR1^-/-^, C5aR2^-/-^ and C5aR1/2^-/-^ mice 18 h after intranasal infection with 2x10^6^ CFU Spn4. (**a**) Principal component analysis (PCA) of the top 500 most variable genes after variance stabilizing transformation of RNA-seq counts (n = 8-10 mice/group). (**b**) Venn diagram showing shared and genotype-specific differentially expressed genes (DEGs). (**c**) Heatmap of normalized expression values of top 50 significant DEGs (baseMean ≥ 1000; Z-score from -2, blue to +2, red). (**d**) Volcano plot showing DEGs in C5aR1/2^-/-^ vs. WT PMN. The number of significantly upregulated and downregulated genes is indicated, with top 15 genes with |log2FC| > 0.58, and C5aR2, labelled. The horizontal dotted line represents adjusted p (adj. p) = 0.05 and vertical lines represent |log2FC| = 1. FC, Fold Change; unadj. p, unadjusted p; ns, not significant. (**e**) Gene ontology (GO) biological processes (BPs) enriched among significant DEGs, highlighting immune-related pathways. pos., positive; neg., negative; reg., regulation. (**f**) Gene set enrichment analysis (GSEA) showing selected immune-related hallmark pathways altered in C5aRs-deficient PMN based on the analysis of significant and non-significant DEGs with Benjamini-Hochberg multiple testing for enrichment. GSEA visualization for C5aR1^-/-^ vs WT PMN depicts processes with adjusted p < 0.2, although adjusted p < 0.05 was used to plot GSEA pathways for C5aR2^-/-^ or C5aR1/2^-/-^ vs WT PMN. NES, normalized enrichment score.

**Extended Data Fig. 7: Single-cell sequencing of BALF leukocytes from WT and C5aR1/2^-/-^ mice.**

Related to Fig. 7. Multimodal sequencing was performed using Cellular Indexing of Transcriptomes and Epitopes by Sequencing (CITE-seq) of FACS-sorted CD45^+^ live BALF cells from WT and C5aR1/2^-/-^ mice (n = 4/group) intranasally infected with 2x10^6^ CFU Spn4 for 18 h. (**a**) Plot showing integrated weighted nearest neighbor uniform manifold approximation and projection (wnnUMAP) overlap in WT and C5aR1/2^-/-^ mice. (**b-c**) Dot plot showing the expression of (**b**) gene and (**c**) protein markers used for cell type annotation. Panel **c** is an expanded plot from **Fig. 7b**. (**d**) Bar plot showing cell type distribution of T, NK, dendritic (DCs), B, and inflammatory monocytes/DC (iMoDCs) cells. pDCs, plasmacytoid DCs; cDCs, classical DCs. (**e**) Violin plot showing expression of interferon genes. (**f**) Violin plot showing module score based on apoptosis related GO BPs across AM clusters. (**g**) Heatmap showing fold change (FC) in multivariate distribution of hallmark gene sets between C5aR1/2^-/-^ and WT mice. Higher/red indicates increase in gene set distribution in C5aR1/2^-/-^ vs. WT. Selected immune-associated gene sets are labelled. *adjusted p < 0.05, **p < 0.01, ***p < 0.001, ****p < 0.0001.

**Extended Data Fig. 8: C5aR1/2^-/-^ mice show differential alteration of AM proliferation and cell death during pneumococcal pneumonia in mice.**

Related to Fig. 8. WT and C5aR1/2^-/-^ mice were intranasally infected with Spn4 (1-2x10^6^ CFU/mouse) or PBS (sham), and BALF were collected 18 h later. (**a, e**) Percentages of live c- Casp-3^+^ or Bax^+^ total Siglec^+^CD11c^+^ alveolar macrophages (AM) were determined by flow cytometry (n ≥ 9/group). (**b, f**) Representative overlay histograms, and (**c, g**) gMFI of cCasp-3 or Bax on AM. Isotype control was included as a reference (n ≥ 8/group). (**d**) Representative flow cytometry plots of Bax on AM from BAL. Cell frequencies of total AM are indicated next to each gate. (**h**) Flow cytometric analysis measuring apoptosis in AM using AnnexinV and Propidium Iodide (PI) staining. Cell frequencies of total AM are indicated next to each gate. (**i**) Bar charts show the percentage of live (AnnexinV^−^PI^−^), apoptotic (AnnexinV^+^PI^-^) and dead (PI^+^) AM in both groups (n ≥ 8/group). (**j**) Representative flow cytometry plots and, (**k**) cell count and fraction of AM that were actively proliferating, expressed as the percentage of total Ki67^+^ AM (n = 8-9/group). (**l**) Representative overlay histograms and (**m**) gMFI of Ki67 on AM. FMO was included as a control (n = 8-9/group). (**n**) Representative flow cytometry dot plots, and (**o**) cell count and fraction of total AM expressing PD-L1 (n = 7-9/group). (**p**) Representative overlay histograms and (**q**) gMFI of PD-L1 expression on PMN. Data are shown as mean ± SEM and are representative of 2-3 independent experiments with similar results. Each circle represents an individual mouse. *p < 0.05, **p < 0.01, ***p < 0.001, ****p < 0.0001, as determined by Student’s t-test. BALF, bronchoalveolar lavage fluid; FMO, fluorescence Minus One; gMFI, geometric mean fluorescence intensity.

## References

1. Van Der Poll, T. & Opal, S.M. Pathogenesis, treatment, and prevention of pneumococcal pneumonia. The Lancet 374, 1543–1556 (2009).

2. Vaughn, V.M., Dickson, R.P., Horowitz, J.K. & Flanders, S.A. Community-Acquired Pneumonia. JAMA (2024).

3. Sirota, S.B. et al. Global burden of lower respiratory infections and aetiologies, 1990– 2023: a systematic analysis for the Global Burden of Disease Study 2023. The Lancet Infectious Diseases 26, 343–361 (2026).

4. Matthay, M.A. et al. Acute respiratory distress syndrome. Nature Reviews Disease Primers 5 (2019).

5. Quinton, L.J., Walkey, A.J. & Mizgerd, J.P. Integrative Physiology of Pneumonia. Physiological Reviews 98, 1417–1464 (2018).

6. Palmer, C.S. & Kimmey, J.M. Neutrophil Recruitment in Pneumococcal Pneumonia. Frontiers in Cellular and Infection Microbiology 12 (2022).

7. Lefrançais, E., Mallavia, B., Zhuo, H., Calfee, C.S. & Looney, M.R. Maladaptive role of neutrophil extracellular traps in pathogen-induced lung injury. JCI Insight 3 (2018).

8. Taenaka, H. et al. Neutrophil Reduction Attenuates the Severity of Lung Injury in the Early Phase of Pneumococcal Pneumonia in Mice. American Journal of Physiology- Lung Cellular and Molecular Physiology (2024).

9. Brown, J.S. et al. The classical pathway is the dominant complement pathway required for innate immunity toStreptococcus pneumoniaeinfection in mice. Proceedings of the National Academy of Sciences 99, 16969–16974 (2002).

10. Gil, E., Noursadeghi, M. & Brown, J.S. Streptococcus pneumoniae interactions with the complement system. Frontiers in Cellular and Infection Microbiology 12 (2022).

11. Jayaraman, A., Walachowski, S. & Bosmann, M. The complement system: A key player in the host response to infections. European Journal of Immunology 54 (2024).

12. Detsika, M.G., Palamaris, K., Dimopoulou, I., Kotanidou, A. & Orfanos, S.E. The complement cascade in lung injury and disease. Respiratory Research 25 (2024).

13. Ricklin, D., Hajishengallis, G., Yang, K. & Lambris, J.D. Complement: a key system for immune surveillance and homeostasis. Nature Immunology 11, 785–797 (2010).

14. Yadav, M.K. et al. Molecular basis of anaphylatoxin binding, activation, and signaling bias at complement receptors. Cell 186, 4956–4973.e4921 (2023).

15. Monk, P.N., Scola, A.M., Madala, P. & Fairlie, D.P. Function, structure and therapeutic potential of complement C5a receptors. British Journal of Pharmacology 152, 429–448 (2007).

16. Gerard, N.P. et al. An Anti-inflammatory Function for the Complement Anaphylatoxin C5a-binding Protein, C5L2. Journal of Biological Chemistry 280, 39677–39680 (2005).

17. Okinaga, S. et al. C5L2, a Nonsignaling C5A Binding Protein. Biochemistry 42, 9406–9415 (2003).

18. Huber-Lang, M. et al. Changes in the Novel Orphan, C5a Receptor (C5L2), during Experimental Sepsis and Sepsis in Humans. The Journal of Immunology 174, 1104–1110 (2005).

19. Bamberg, C.E. et al. The C5a Receptor (C5aR) C5L2 Is a Modulator of C5aR-mediated Signal Transduction. Journal of Biological Chemistry 285, 7633–7644 (2010).

20. Chen, N.-J. et al. C5L2 is critical for the biological activities of the anaphylatoxins C5a and C3a. Nature 446, 203–207 (2007).

21. Yu, S. et al. The complement receptor C5aR2 promotes protein kinase R expression and contributes to NLRP3 inflammasome activation and HMGB1 release from macrophages. Journal of Biological Chemistry 294, 8384–8394 (2019).

22. Wang, R., Lu, B., Gerard, C. & Gerard, N.P. Disruption of the Complement Anaphylatoxin Receptor C5L2 Exacerbates Inflammation in Allergic Contact Dermatitis. The Journal of Immunology 191, 4001–4009 (2013).

23. Wang, R., Lu, B., Gerard, C. & Gerard, N.P. C5L2, the Second C5a Anaphylatoxin Receptor, Suppresses LPS-Induced Acute Lung Injury. American Journal of Respiratory Cell and Molecular Biology 55, 657–666 (2016).

24. Wu, M.C.L., Lee, J.D., Ruitenberg, M.J. & Woodruff, T.M. Absence of the C5a Receptor C5aR2 Worsens Ischemic Tissue Injury by Increasing C5aR1-Mediated Neutrophil Infiltration. The Journal of Immunology 205, 2834–2839 (2020).

25. Woehrl, B. et al. Complement component 5 contributes to poor disease outcome in humans and mice with pneumococcal meningitis. Journal of Clinical Investigation 121, 3943–3953 (2011).

26. Lu, Z., Zhu, L., Yi, C., Su, B. & Wang, R. C5a/C5aR regulates Th1/Th2 imbalance in sepsis-associated lung injury by promoting neutrophil activation to increase PAD4 expression. Annals of Medicine 57 (2025).

27. Kambas, K. et al. C5a and TNF-α Up-Regulate the Expression of Tissue Factor in Intra- Alveolar Neutrophils of Patients with the Acute Respiratory Distress Syndrome. The Journal of Immunology 180, 7368–7375 (2008).

28. Sadik, C.D., Kim, N.D., Iwakura, Y. & Luster, A.D. Neutrophils orchestrate their own recruitment in murine arthritis through C5aR and FcγR signaling. Proceedings of the National Academy of Sciences 109, E3177–E3185 (2012).

29. Leonardi, L. et al. Inherited defects in the complement system. Pediatric Allergy and Immunology 33, 73–76 (2022).

30. Russkamp, N.F. et al. Experimental design of complement component 5a-induced acute lung injury (C5a-ALI): a role of CC-chemokine receptor type 5 during immune activation by anaphylatoxin. The FASEB Journal 29, 3762–3772 (2015).

31. Chaudhary, N., Jayaraman, A., Reinhardt, C., Campbell, J.D. & Bosmann, M. A single- cell lung atlas of complement genes identifies the mesothelium and epithelium as prominent sources of extrahepatic complement proteins. Mucosal Immunology 15, 927–939 (2022).

32. Tung, C.-H. et al. In Vivo Imaging of β-Galactosidase Activity Using Far Red Fluorescent Switch. Cancer Research 64, 1579–1583 (2004).

33. Sandgren, A. et al. Virulence in Mice of Pneumococcal Clonal Types with Known Invasive Disease Potential in Humans. The Journal of Infectious Diseases 192, 791–800 (2005).

34. Yousefi, S., Mihalache, C., Kozlowski, E., Schmid, I. & Simon, H.U. Viable neutrophils release mitochondrial DNA to form neutrophil extracellular traps. Cell Death & Differentiation 16, 1438–1444 (2009).

35. Bosmann, M. et al. Extracellular histones are essential effectors of C5aR- and C5L2- mediated tissue damage and inflammation in acute lung injury. FASEB J 27, 5010–5021 (2013).

36. Lin, R., Zhang, Y., Pradhan, K. & Li, L. TICAM2-related pathway mediates neutrophil exhaustion. Scientific Reports 10 (2020).

37. Li, R., Coulthard, L.G., Wu, M.C.L., Taylor, S.M. & Woodruff, T.M. C5L2: a controversial receptor of complement anaphylatoxin, C5a. The FASEB Journal 27, 855–864 (2013).

38. Rittirsch, D. et al. Functional roles for C5a receptors in sepsis. Nature Medicine 14, 551–557 (2008).

39. Ward, P.A. The Harmful Role of C5a on Innate Immunity in Sepsis. Journal of Innate Immunity 2, 439–445 (2010).

40. Hajishengallis, G. & Lambris, J.D. Crosstalk pathways between Toll-like receptors and the complement system. Trends in Immunology 31, 154–163 (2010).

41. Li, X.X., Lee, J.D., Kemper, C. & Woodruff, T.M. The Complement Receptor C5aR2: A Powerful Modulator of Innate and Adaptive Immunity. J Immunol 202, 3339–3348 (2019).

42. Hsu, W.-C., Yang, F.-C., Lin, C.-H., Hsieh, S.-L. & Chen, N.-J. C5L2 is required for C5a-triggered receptor internalization and ERK signaling. Cellular Signalling 26, 1409–1419 (2014).

43. Croker, D.E. et al. C5a2 can modulate ERK1/2 signaling in macrophages via heteromer formation with C5a1 and β-arrestin recruitment. Immunology & Cell Biology 92, 631–639 (2014).

44. Brinkmann, V. et al. Neutrophil Extracellular Traps Kill Bacteria. Science 303, 1532–1535 (2004).

45. Park, S.Y. et al. Autophagy Primes Neutrophils for Neutrophil Extracellular Trap Formation during Sepsis. American Journal of Respiratory and Critical Care Medicine 196, 577–589 (2017).

46. Pilsczek, F.H. et al. A Novel Mechanism of Rapid Nuclear Neutrophil Extracellular Trap Formation in Response to Staphylococcus aureus. The Journal of Immunology 185, 7413–7425 (2010).

47. Yipp, B.G. et al. Infection-induced NETosis is a dynamic process involving neutrophil multitasking in vivo. Nature Medicine 18, 1386–1393 (2012).

48. Shen, X., Cao, K., Zhao, Y. & Du, J. Targeting Neutrophils in Sepsis: From Mechanism to Translation. Frontiers in Pharmacology 12 (2021).

49. Urban, C. & Zychlinsky, A. Netting bacteria in sepsis. Nature Medicine 13, 403–404 (2007).

50. Tuz, A.A. et al. Stroke and myocardial infarction induce neutrophil extracellular trap release disrupting lymphoid organ structure and immunoglobulin secretion. Nature Cardiovascular Research 3, 525–540 (2024).

51. Guan, Z., et al. Impaired AMP-Dependent Protein Kinase-Mediated Neutrophil Extracellular Trap Clearance by Aged Macrophages in Sepsis-Induced Liver Injury. Anesthesia & Analgesia (2025).

52. Poli, V. & Zanoni, I. Neutrophil intrinsic and extrinsic regulation of NETosis in health and disease. Trends in Microbiology 31, 280–293 (2023).

53. Sanders, N.L. et al. Neutrophil Extracellular Traps as an Exacerbating Factor in Bacterial Pneumonia. Infect Immun 90, e0049121 (2022).

54. Xue, J., Ren, Y., Chen, Y., Zhu, H. & Wang, Y. Inhibition of neutrophil infiltration and NETosis ameliorates sepsis-induced cardiac injury and reduces myocardial inflammation and apoptosis. International Immunopharmacology 168, 115862 (2026).

55. Yipp, B.G. & Kubes, P. NETosis: how vital is it? Blood 122, 2784–2794 (2013).

56. Zhang, J. et al. Dysregulation of neutrophil in sepsis: recent insights and advances. Cell Communication and Signaling 23 (2025).

57. Farrera, C. & Fadeel, B. Macrophage Clearance of Neutrophil Extracellular Traps Is a Silent Process. The Journal of Immunology 191, 2647–2656 (2013).

58. Grégoire, M. et al. Impaired efferocytosis and neutrophil extracellular trap clearance by macrophages in ARDS. European Respiratory Journal 52, 1702590 (2018).

59. Shan, X. et al. Interactions between neutrophil extracellular traps and macrophages: the key to inflammatory diseases. Frontiers in Immunology 17 (2026).

60. Monteith, A.J., Miller, J.M., Maxwell, C.N., Chazin, W.J. & Skaar, E.P. Neutrophil extracellular traps enhance macrophage killing of bacterial pathogens. Science Advances 7 (2021).

61. Apel, F. et al. The cytosolic DNA sensor cGAS recognizes neutrophil extracellular traps. Science Signaling 14, eaax7942 (2021).

62. Alves-Filho, J.C., Spiller, F. & Cunha, F.Q. NEUTROPHIL PARALYSIS IN SEPSIS. Shock 34, 15–21 (2010).

63. Palomino-Segura, M., Sicilia, J., Ballesteros, I. & Hidalgo, A. Strategies of neutrophil diversification. Nature Immunology 24, 575–584 (2023).

64. Schulte-Schrepping, J. et al. Severe COVID-19 Is Marked by a Dysregulated Myeloid Cell Compartment. Cell 182, 1419–1440.e1423 (2020).

65. Ma, C., Wang, L., Wang, X., Zhu, X. & Chen, Y. PD-L1+ neutrophils trigger pulmonary endothelial pyroptosis via an oxidative phosphorylation-dependent mechanism in sepsis. Respiratory Research 26 (2025).

66. Martelossi Cebinelli, G.C., et al. CXCR4+ PD-L1+ neutrophils are increased in non- survived septic mice. iScience 28, 112083 (2025).

67. Thanabalasuriar, A. et al. PD-L1 + neutrophils contribute to injury-induced infection susceptibility. Science Advances 7, eabd9436 (2021).

68. Wang, J.-F. et al. Up-regulation of Programmed Cell Death 1 Ligand 1 on Neutrophils May Be Involved in Sepsis-induced Immunosuppression. Anesthesiology 122, 852–863 (2015).

69. De Kleijn, S., et al. IFN-γ-Stimulated Neutrophils Suppress Lymphocyte Proliferation through Expression of PD-L1. PLoS ONE 8, e72249 (2013).

70. Langereis, J.D. et al. Spleen-derived IFN-γ induces generation of PD-L1+-suppressive neutrophils during endotoxemia. Journal of Leukocyte Biology 102, 1401–1409 (2017).

71. Wang, J.-F. et al. Upregulated PD-L1 delays human neutrophil apoptosis and promotes lung injury in an experimental mouse model of sepsis. Blood 138, 806–810 (2021).

72. Garcia-Diaz, A. et al. Interferon Receptor Signaling Pathways Regulating PD-L1 and PD-L2 Expression. Cell Reports 19, 1189–1201 (2017).

73. Zhu, C.-L. et al. PD-L1 maintains neutrophil extracellular traps release by inhibiting neutrophil autophagy in endotoxin-induced lung injury. Frontiers in Immunology 13 (2022).

74. Müller-Redetzky, H. et al. Neutralizing Complement C5a Protects Mice with Pneumococcal Pulmonary Sepsis. Anesthesiology 132, 795–807 (2020).

75. Huber-Lang, M.S., et al. Protective effects of anti-C5a peptide antibodies in experimental sepsis. The FASEB Journal 15, 568–570 (2001).

76. Panas, R.M., Burnett, B.P., Chong, C., Guo, R. & Riedemann, N. Treatment of Viral- induced Acute Respiratory Distress Syndrome (ARDS) With Vilobelimab: A Focus on C5a Inhibition. American Journal of Respiratory and Critical Care Medicine 211, A3110–A3110 (2025).

77. Okusa, S. et al. Serious Bacterial Infections Associated with Eculizumab: A Pharmacovigilance Study. Internal Medicine 63, 1061–1066 (2024).

## Methods References

78. Concordet, J.-P. & Haeussler, M. CRISPOR: intuitive guide selection for CRISPR/Cas9 genome editing experiments and screens. Nucleic Acids Research 46, W242–W245 (2018).

79. Tettelin, H. et al. Complete genome sequence of a virulent isolate of Streptococcus pneumoniae. SCIENCE 293, 498–506 (2001).

80. Dudek, M. et al. Lung epithelium and myeloid cells cooperate to clear acute pneumococcal infection. Mucosal Immunol 9, 1288–1302 (2016).

81. Stagg, A.J., Burke, F., Hill, S. & Knight, S.C. Isolation of mouse spleen dendritic cells. Methods Mol Med 64, 9–22 (2001).

82. Cossarizza, A. et al. Guidelines for the use of flow cytometry and cell sorting in immunological studies (third edition). Eur J Immunol 51, 2708–3145 (2021).

83. Arafa, E.I., et al. Recruitment and training of alveolar macrophages after pneumococcal pneumonia. JCI Insight 7 (2022).

84. Rue-Albrecht, K., Marini, F., Soneson, C. & Lun, A.T.L. iSEE: Interactive SummarizedExperiment Explorer. F1000Research 7, 741 (2018).

