## Supplemental Material for "Dual C5aR1 and C5aR2 deficiency uncouples neutrophilic inflammation from antibacterial host defense"

### Supplementary Methods

#### *Whole Genome sequencing*

To exclude off target mutations in the CRISPR-generated C5aR1/2$^{-/-}$ mouse colony, lung genomic DNA (gDNA) collected from a C5aR1/2$^{-/-}$ naive mouse was extracted using the QIAamp Fast DNA Tissue kit (Qiagen) following the manufacturer's recommendations. The gDNA was quantified using a Nanodrop 2000c instrument and integrity was assessed by agarose gel electrophoresis. All gDNA samples showed a single intact band at 23 kb without any sign of smearing. The whole genome was sequenced on the Illumina NovaSeq 6000 platform. Adapters were trimmed using Cutadapt (v. 2.10)[1]. The Genome Analysis Toolkit (GATK,v. 4.1.2.0) preprocessing protocols were used to process the raw reads[2]. The reads were mapped to the *Mus musculus* reference genome GRCm38 from the Ensembl database (ftp://ftp.ensembl.org/pub/release-101/) using the BWA mem algorithm[3]. The PCR duplicates were removed using Picard (v. 2.18.11) (http://broadinstitute.github.io/picard/). Variant calling and filtration for single nucleotide polymorphisms (SNPs) and small Indels was performed using GATK HaplotypeCaller and VariantFiltration. The hard filtering parameters for single nucleotide variants (SNVs) were set as follows: DP < 10, QD < 2.0, QUAL < 30.0, SOR > 3.0, FS > 60.0, MQ < 40.0, MQRankSum ≤ 12.5, ReadPosRankSum ≤ 8.0 and for INDELS were: DP < 10, QD < 2.0, FS > 200.0, MQ < 40.0, ReadPosRankSum ≤ 20.0. Large deletions were identified using structural variant callers SvABA[4] (v. 1.1.0) and DELLY[5] (v. 0.7.9). The predicted off-target regions in the reference genome were identified using CRISPOR (v. 4.98)[6], Cas-OFFinder[7] and BWA aln algorithms[3]. For Cas-OFFinder and BWA aln, either up to four mismatches or a single DNA/RNA bulge together with up to three mismatches relative to the sgRNA sequence were allowed, excluding the PAM sequence. Off-target sequences were considered only in the presence of downstream NGG, NAG or NGA PAM motifs. There were 5298 unique off-target sites in C5aR1/2 knockout mice. The variants in the samples were compared to the locations of the predicted off-target regions. Any variant within 20 nucleotides up-/downstream of the predicted off-target site was manually curated. Any variants documented in the ensemble database (ftp://ftp.ensembl.org/pub/release-101/variation/vcf/) were filtered out. Large deletions (> 50 nucleotides) commonly identified by SvABA and DELLY were also manually analyzed irrespective of their genomic location.

The whole genome sequence of the CRISPR/Cas9 C5aR1/2$^{-/-}$ mouse was analyzed for SNPs and Indels located 20 nucleotides up-/downstream of the gRNA homologous sites to exclude off-target effects. No variants were identified around the off-target sites. Deletions larger than 50 nucleotides present across the genome were validated manually and were not a result of CRISPR mutagenesis.

#### *Bulk RNA-seq library preparation and data analysis*

BALF cells were prepared following the same protocol as for flow cytometry. Alveolar macrophages (AM) were defined as FVD$^-$CD64$^+$Ly6G$^-$SiglecF$^+$CD11c$^+$ singlets and FVD$^-$CD64$^-$Ly6G$^+$SiglecF$^-$CD11c$^-$ singlets as neutrophils (PMN) during FACSorting (BD FACSAria™ II SORP, BD Biosciences). Total RNA was extracted from those populations using the Qiagen RNeasy Mini Kit (QIAGEN, Germantown, USA) or InnuPREP RNA Mini kit (Analytik Jena, Berlin, Germany). RNA abundance was quantified using a Qubit RNA HS Assay kit (Life Technologies, Thermo Fisher Scientific) on a Qubit fluorometer (Life Technologies). RNA integrity was evaluated spectrophotometrically with RNA 6000 Nano or Pico total RNA kits (Agilent, Santa Clara, USA) on an Agilent 2100 Bioanalyzer (Agilent Technologies). All samples showed a RIN value of ≥8. The RNA-seq libraries were constructed with NEBNext Low-Input RNA Library Prep Kit for Illumina with an extra SMARTer RNA amplification step for neutrophils and sequenced with paired-end reads on a PE150 Illumina NovaSeq 6000 (Novogene, Yuen Long, Hong Kong).

The bulk RNA-seq data (C5a-ALI, C5aR1$^{-/-}$, C5aR2$^{-/-}$, C5aR1/2$^{-/-}$ AM and PMN) were analyzed using the Boston University Shared Computing Cluster and a desktop computer running Linux Mint (v. 19.3) following standard methodology[8]. The paired-end FASTQ files were aligned to the reference mouse genome (GENCODE Release M25, GRCm38.p6 [C5a-ALI]; Ensembl GRCm38.76 [C5aR1$^{-/-}$, C5aR2$^{-/-}$, C5aR1/2$^{-/-}$ AM, PMN]) with STAR RNA-seq aligner (v. 2.6.1a /2.7.10a)[9] and indexed using the samtools (v. 1.10)[10] software suite. The featureCounts tool from the subread/Rsubread package (v. 1.6.2/v. 2.0.1)[11] was used to quantify the reads. The sequenced reads and counts matrix were quality checked using the FastQC (v. 0.11.7) and MultiQC (v. 1.6)[12] tools. Exploratory and differential expression analyses were performed with the DESeq2 package (v. 1.28.1)[13] through RStudio Server (v. 1.3.1073) running the R framework (4.0.2 and 4.0.5). Outlier samples detected in each experiment (n = 1/group) which showed low sequence quality were excluded from the analysis. Differential expression analysis with DESeq2 was based on Wald's test with Benjamini-Hochberg multiple testing method. Shrinkage of large log2 fold change values was done using the apeglm (v. 1.23.1) package (C5a-ALI)[14] and the threshold for statistical significance set at adjusted $p < 0.05$. Gene ontology biological process (GO BP) enrichment analyses was performed using the topGO (v. 2.40.0)[15] and GeneTonic (v.

1.0.1/1.5.2)[14] packages, while the clusterProfiler (v. 3.16.0)[16] package was used for Kyoto Encyclopedia of Genes and Genomes (KEGG) pathway enrichment analysis. Both analyses were performed with statistically significant differentially expressed genes with adjusted $p < 0.05$, with the background for enrichment set to all expressed genes. Hallmark gene sets for mouse were retrieved with the msigdbr (v. 7.4.1) package for gene set enrichment analysis (GSEA) using clusterProfiler with Benjamini-Hochberg multiple testing (C5aR1$^{-/-}$, C5aR2$^{-/-}$, C5aR1/2$^{-/-}$ AM, PMN)[17]. The GSEA included both significant and non-significant differentially expressed genes, ranked based on the product of -log10(unadjusted p value) and sign(log2 fold change); p values close to zero due to detection limits were set to $10^{-1}$ x minimum(unadjusted p value) to circumvent undefined logarithmic conversion with zero values[18]. The statistical significance threshold was set at p.value elim < 0.05 for topGO GO BP and adjusted $p < 0.05$ for KEGG and GSEA. Only for C5aR1$^{-/-}$ vs WT PMN GSEA, the significance threshold was set at adjusted $p < 0.2$, to broaden the search for immune associated geneset enrichment. Normalized enrichment score (NES) < 0 indicates downregulation of hallmark genesets whereas NES > 0 indicates enrichment for the corresponding processes. The enriched genes were visualized through heatmaps generated from the pheatmap (v. 1.0.12)[19] and Complex Heatmap (v. 2.4.3)[20] packages. All other visualizations were based on in-built functions from the DESeq2, ggplot2 (v. 3.3.2)[21], VennDetail (v. 1.4.0)[22], clusterProfiler, and pcaExplorer (v. 2.14.2)[23] packages.

#### *Single-cell RNA-seq (CITE-seq/Total-seq)*

*Sample preparation*

We used BALF cells from WT and C5aR1/2$^{-/-}$ mice (n = 4 mice/group) after intranasal infection with $2x10^6$ CFU of Spn TIGR4 for 18 h. Following incubation in Fc receptor blocking reagent (TruStain FcX™ PLUS, BioLegend) diluted in 1X MACS Buffer (2.5 mM EDTA and 1% BSA in PBS) for 10 min at 4°C, cells were stained with FVD and anti-CD45 antibody for 20 min at 4°C. Singlets that were FVD$^-$CD45$^+$ were sorted using an BD FACSAria™ II SORP cell sorter (BD Biosciences). Next, FACS-sorted leukocytes were incubated with a mouse TotalSeq™-A pretitrated cocktail of 198 antibodies (custom panel, BioLegend) listed in **Supplementary Table 5** for 30 min at 4°C according to the manufacturer's specifications. An additional TotalSeq™-A 'Spike-in' antibody (Totalseq™-A1042 anti-mouse CD88, clone 20/70, BioLegend) was added to the staining panel to target C5aR1 protein following an optimized titration protocol. After staining, the cells were washed 3 times by resuspension in 1X CITE-seq buffer (0.04% BSA in PBS) followed by centrifugation (650 g for 10 min at 4°C) and supernatant disposal. Cell viability was assessed for each sample using Trypan Blue (1:1) in a Neubauer hematocytometer (Hausser Scientific), and the few samples that fell below 90% viability were not processed further.

*10X processing and library preparation*

A target of ~3,000 cells per sample was captured using a Chromium Next GEM single-cell 3' Gel Bead kit v3.1 on each well of a Chip G (10X Genomics). The preparation of antibody-derived Tag (ADT)- and mRNA-derived cDNA libraries from reverse transcription to final amplification, was performed with a KAPA HiFi Hotstart ReadyMix PCR kit (KAPA Biosystems) and a Chromium Next GEM single-cell 3′ GEM library kit v3.1 (10X Genomics), respectively. For ADT libraries, additional primers were included at the cDNA amplification step (ADT Additive Primer) and the library construction steps (SI PCR primer and Small RNA RPI1 to RPI8 primers, index i7) as recommended by BioLegend. Quality control of generated intermediate libraries was performed using a High Sensitivity DNA chip run on a Bioanalyzer 2100 device (Agilent). ADT libraries contained a predominant single peak at 180 bp whereas mRNA-derived cDNA libraries were featured by a single peak at ~500 bp.

The mRNA-derived cDNA libraries were indexed using a Single Index kit (Single Index T set A,10X Genomics) and further combined with ADT libraries at desired proportions (10% and 90% of a lane, respectively) for each sample. Pooled Gene Expression (mRNA-derived cDNA) and Cell Surface Protein (ADT) libraries were paired-end sequenced on a NextSeq2000 device (Illumina) at a depth of 1.33 billion reads (92% > 30 and 0.12% error rate), resulting in a read depth of 55,000 reads per cell on average (5,000 reads per cells for ADTs and 50,000 reads per cell for mRNA-derived cDNAs).

*Raw data analysis*

The CITE-Seq/TotalSeq data of the 8 samples (WT and C5aR1/2$^{-/-}$; n = 4 each) were analyzed using the Boston University Shared Computing Cluster resources. The Illumina base call files were demultiplexed using the 10X Genomics Cell Ranger (v. 6.0.1) mkfastq module. The demultiplexed FASTQ files were aligned using pre-built mouse reference from 10X Genomics (mm10; GENCODE vM23/Ensembl 98) and quantified using Cell Ranger's count module. The raw counts matrix was imported into RStudio Server (v. 1.3.1073) running the R framework (v.

4.1.1) for analysis using the Seurat package (v. 4.0.4)[24]. During the quality control step, sample-specific filtering was performed to only retain cells having ≥ 200 and ≤ 6000/6500 genes/cell, ≤ 40,000 RNA Unique Molecular Identifier (UMI) counts/cell, ≤ 4000/5000/6000 ADT counts/cell and percent mitochondrial reads ≤ 10%. Doublet identification was performed using the scDblFinder (v. 1.8.0)[25] package based on an estimated doublet proportion of 2.41% (calculated from the Satija Lab's multiplexing cost calculator: https://satijalab.org/costpercell/). Briefly, scDblFinder uses a classifier built on gradient boosted trees based on predictors such as library size and marker coexpression to differentiate "singlets" and "doublets". Cells identified as doublets were filtered out. As a cross-validation, we also checked that the retained singlets do not express markers of more than one cell type, a characteristic of doublet/multiplets.

Each sample was normalized using Seurat's SCTransform function for the RNA assay and a centered log-ratio (CLR) transformation for the ADT assay. Integration of the samples was performed separately for each assay following Seurat's standard integration workflow. The integrated dataset was checked for segregation based on cell cycle phase. The integrated ADT assay data was added to the Seurat object containing the integrated RNA data following the standard Seurat workflow recommendations. This step was followed by dimensional reduction using the Principal Component Analysis (PCA) and Uniform Manifold Approximation and Projection (UMAP) methods for the RNA and ADT assays. Seurat's weighted-nearest neighbor (WNN) approach was used to identify multimodal neighbors based on the first 30 principal components (PCs) from RNA and the first 20 PCs from ADT. The cells were clustered using the smart local moving (SLM) algorithm over a range of resolutions from 0.5–4 based on the WNN graph. After assessment of cluster stability using the clustree (v. 0.4.4)[26] package and the silhouette coefficient from the cluster package (v. 2.1.2)[27], resolution 1.1 was selected for further analysis. Cell types were identified based on gene and protein markers from literature. Sub-clustering was performed to resolve the identities of B cell, conventional dendritic cell (cDC) and plasmacytoid dendritic cell (pDC) clusters, and $CD4^+$, $CD8^+$ and γδ T cell clusters.

Differential expression (DE) analysis was performed separately for the RNA and ADT assays between WT and $C5aR1/2^{-/-}$ genotypes for each cell type using Seurat's FindMarkers function with the Wilcoxon Rank-Sum test and multiple hypothesis testing using Bonferroni correction method. The logfc.threshold parameter was set to the default threshold of 0.25 for log2 fold-change (log2FC) for the RNA and ADT DE analysis. The min.pct was set to the default of 0.1 to only test the genes and ADTs that were expressed in more than 10% of the cells for both analyses. GO BP enrichment analysis on the DE genes for each cell type was performed using topGO through the pcaExplorer (v. 2.18.0)[23] package, with the background set to genes expressed in at least one cell of the respective cell type. Visualizations were performed using Seurat's visualization functions and the ggplot2 (v. 3.3.5)[21] package. Differential abundance analysis comparing cell type proportions between C5aR1/2$^{-/-}$ and WT was performed using edgeR's (v3.34.1) quasi-likelihood negative binomial generalized linear model and generalized linear model likelihood ratio test[28, 29]. Differential abundance analysis comparing cell type proportions between C5aR1/2$^{-/-}$ and WT was performed using edgeR's (v. 3.34.1) quasi-likelihood negative binomial generalized linear model and generalized linear model likelihood ratio, with Benjamini-Hochberg multiple testing[30]. Module scores for apoptosis enriched genes from the GO BP results for neutrophil and alveolar macrophage clusters were estimated using Seurat's AddModuleScore function, and statistical significance calculated using the Wilcox_test with Benjamini-Hochberg multiple testing in the rstatix package (v. 0.7.0)[31]. A systems level characterization of hallmark genesets across the cell types between C5aR1/2$^{-/-}$ and WT mice was performed using the single-cell pathway analysis (SCPA; v. 1.6.2) package[32]. Hallmark gene sets for *Mus musculus* were retrieved using the msigdbr package (v. 7.5.1). SCPA estimates the change in multivariate distribution in pathways/gene sets between groups for each cell type and outputs both an enrichment metric (fold change) and an estimate of change in multivariate distribution (q value). Larger q values indicate greater change in pathway/gene set activity. SCPA enrichment results were visualized using ComplexHeatmap. The SCPA estimations were performed in R v. 4.3.2. The statistical significance threshold for DE, and cell type abundance, and module analyses were set at adjusted $p < 0.05$, and for GO BP at p.value elim < 0.05.

### References


1. Martin, M. Cutadapt removes adapter sequences from high-throughput sequencing reads. *EMBnet.journal* **17**, 10 (2011).

2. Mckenna, A. *et al.* The Genome Analysis Toolkit: A MapReduce framework for analyzing next-generation DNA sequencing data. *Genome Research* **20**, 1297-1303 (2010).

3. Li, H. & Durbin, R. Fast and accurate short read alignment with Burrows-Wheeler transform. *Bioinformatics* **25**, 1754-1760 (2009).

4. Wala, J.A. *et al.* SvABA: genome-wide detection of structural variants and indels by local assembly. *Genome Research* **28**, 581-591 (2018).

5. Rausch, T. *et al.* DELLY: structural variant discovery by integrated paired-end and split-read analysis. *Bioinformatics* **28**, i333-i339 (2012).

6. Concordet, J.-P. & Haeussler, M. CRISPOR: intuitive guide selection for CRISPR/Cas9 genome editing experiments and screens. *Nucleic Acids Research* **46**, W242-W245 (2018).

7. Bae, S., Park, J. & Kim, J.S. Cas-OFFinder: a fast and versatile algorithm that searches for potential off-target sites of Cas9 RNA-guided endonucleases. *Bioinformatics* **30**, 1473-1475 (2014).

8. Ludt, A., Ustjanzew, A., Binder, H., Strauch, K. & Marini, F. Interactive and Reproducible Workflows for Exploring and Modeling RNA-seq Data with pcaExplorer, Ideal, and GeneTonic. *Curr Protoc* **2**, e411 (2022).

9. Dobin, A. *et al.* STAR: ultrafast universal RNA-seq aligner. *Bioinformatics* **29**, 15-21 (2013).

10. Danecek, P. *et al.* Twelve years of SAMtools and BCFtools. *Gigascience* **10** (2021).

11. Liao, Y., Smyth, G.K. & Shi, W. featureCounts: an efficient general purpose program for assigning sequence reads to genomic features. *Bioinformatics* **30**, 923-930 (2014).

12. Ewels, P., Magnusson, M., Lundin, S. & Käller, M. MultiQC: summarize analysis results for multiple tools and samples in a single report. *Bioinformatics* **32**, 3047-3048 (2016).

13. Love, M.I., Huber, W. & Anders, S. Moderated estimation of fold change and dispersion for RNA-seq data with DESeq2. *Genome Biology* **15** (2014).

14. Marini, F., Ludt, A., Linke, J. & Strauch, K. GeneTonic: an R/Bioconductor package for streamlining the interpretation of RNA-seq data. *BMC Bioinformatics* **22**, 610 (2021).

15. Alexa, A. & Rahnenfuhrer, J. topGO: Enrichment Analysis for Gene Ontology. R package version 2.40.0. 2021.

16. Yu, G., Wang, L.G., Han, Y. & He, Q.Y. clusterProfiler: an R package for comparing biological themes among gene clusters. *Omics* **16**, 284-287 (2012).

17. Dolgalev, I. msigdbr: MSigDB Gene Sets for Multiple Organisms in a Tidy Data Format. R package version 7.4.1. https://CRAN.R-project.org/package=msigdbr. 2021.

18. Blighe, K., Rana, S. & Lewis, M. EnhancedVolcano: Publication-ready volcano plots with enhanced colouring and labeling. R package version 1.14.0, https://github.com/kevinblighe/EnhancedVolcano. 2022.

19. Kolde, R. pheatmap: Pretty Heatmaps. R package version 1.0.12. 2019.

20. Gu, Z., Eils, R. & Schlesner, M. Complex heatmaps reveal patterns and correlations in multidimensional genomic data. *Bioinformatics* **32**, 2847-2849 (2016).

21. Wickham, H. *ggplot2: Elegant Graphics for Data Analysis*, 2 edn. Springer International Publishing, 2016.

22. Guo, K. & McGregor, B. VennDetail: A package for visualization and extract details. R package version 1.12.0, https://github.com/guokai8/VennDetail. 2022.

23. Marini, F. & Binder, H. pcaExplorer: an R/Bioconductor package for interacting with RNA-seq principal components. *BMC Bioinformatics* **20**, 331 (2019).

24. Hao, Y. *et al.* Integrated analysis of multimodal single-cell data. *Cell* **184**, 3573-3587.e3529 (2021).

25. Germain, P., Lun, A., Garcia Meixide, C., Macnair, W. & Robinson, M. Doublet identification in single-cell sequencing data using scDblFinder [version 2; peer review: 2 approved]. *F1000Research* **10** (2022).

26. Zappia, L. & Oshlack, A. Clustering trees: a visualization for evaluating clusterings at multiple resolutions. *Gigascience* **7**, giy083 (2018).

27. Maechler, M., Rousseeuw, P., Struyf, A., Hubert, M. & Hornik, K. cluster: Cluster Analysis Basics and Extensions. R package version 2.1.2. 2021.

28. Crowell, H.L. *et al.* muscat detects subpopulation-specific state transitions from multi-sample multi-condition single-cell transcriptomics data. *Nat Commun* **11**, 6077 (2020).

29. Robinson, M.D., McCarthy, D.J. & Smyth, G.K. edgeR: a Bioconductor package for differential expression analysis of digital gene expression data. *Bioinformatics* **26**, 139-140 (2010).

30. Walter, W., Sánchez-Cabo, F. & Ricote, M. GOplot: an R package for visually combining expression data with functional analysis. *Bioinformatics* **31**, 2912-2914 (2015).

31. Kassambara, A. rstatix: Pipe-Friendly Framework for Basic Statistical Tests. R package version 0.7.0. https://CRAN.R-project.org/package=rstatix. 2021.

32. Bibby, J.A. *et al.* Systematic single-cell pathway analysis to characterize early T cell activation. *Cell Rep* **41**, 111697 (2022).

33. Rue-Albrecht, K., Marini, F., Soneson, C. & Lun, A.T.L. iSEE: Interactive SummarizedExperiment Explorer. *F1000Research* **7**, 741 (2018).

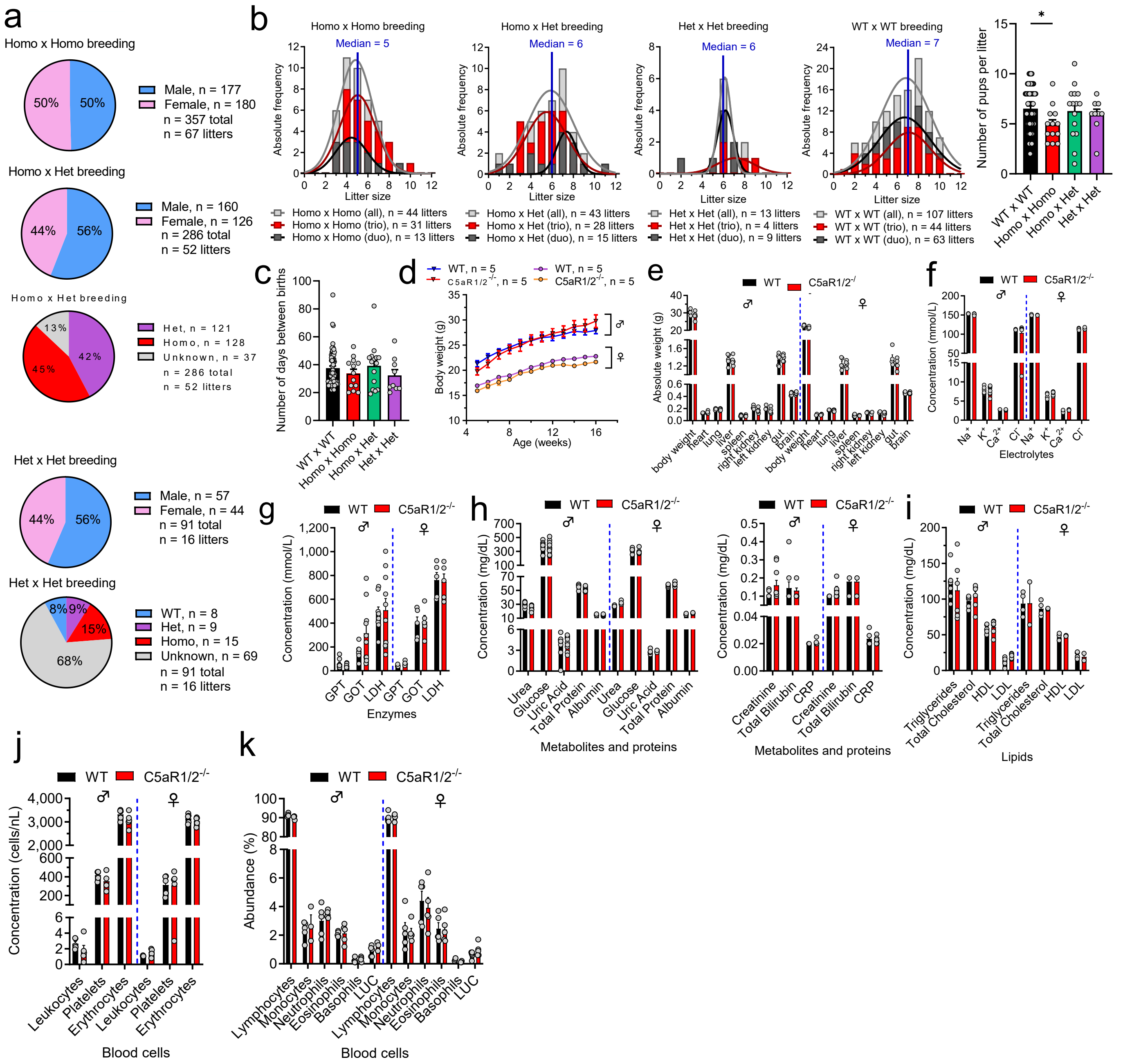

**Extended Data Fig.1: Breeding statistics and blood analysis of C5aR1/2$^{-/-}$ double knockout mice.** Related to Fig. 1. (**a-c**) Analysis of breeding data collected over a 24-month period from inbred mice of C5aR1/2$^{-/-}$ and C57BL/6J wild type (WT) strains. (**a**) Frequency of genders and genotypes of mice born in each configuration of breeding cages. The upper panel depicts the sex ratio of pups born from homozygous (Homo x Homo) breeders, the two charts below show the genders and genotypes distribution across pups born from the breeding of heterozygous and homozygous (Het x Homo) mice, and the lowest two panels represent similar distribution in heterozygous breeding pairs (Het x Het). The number of litters and offspring corresponding to each breeding condition is indicated in the legend of each chart. Unknown genotype frequencies represent offspring mice that were not sampled for genotyping. (**b-c**) Analysis of reproductive performance metrics of pair-bred (duo) and trio-bred (trio) mice for each category of breeding condition. (**b**) Bar graphs represent the absolute frequencies (number of litters) across litter size categories. The median of litter size is presented in blue and the fit curves show the frequency distribution of litter size for each breeding configuration. The corresponding total number of litters is stated below each graph. The right panel summarizes the average litter size in each duo-bred cage. (**c**) Number of days between births in each breeding configuration. Values are represented as mean ± SEM. (**d-k**) C5aR1/2$^{-/-}$ mice (females: ♀; males: ♂) and their C57BL/6J (WT) counterparts were (**d**) weighted weekly from 4-16 weeks after birth. (**e**) Mice were necropsied at 17 weeks of age and organ weights were recorded. (**f-h**) Clinical chemistry analysis of peripheral blood. (**i**) Quantification of lipids. (**j-k**) Differential blood cell count. Bar graphs show mean ± SEM values from the analysis of n = 5 mice per strain and per gender. Data are representative of 3 independent experiments. *$p < 0.05$ as determined by Student's t-test analysis. GPT, Alanine aminotransferase; GOT, Aspartate aminotransferase; LDH, Lactate Dehydrogenase; CRP, C Protein Reactive; HDL, High Density Lipoprotein; LDL, Low Density Lipoprotein; LUC, Large unstained cells.

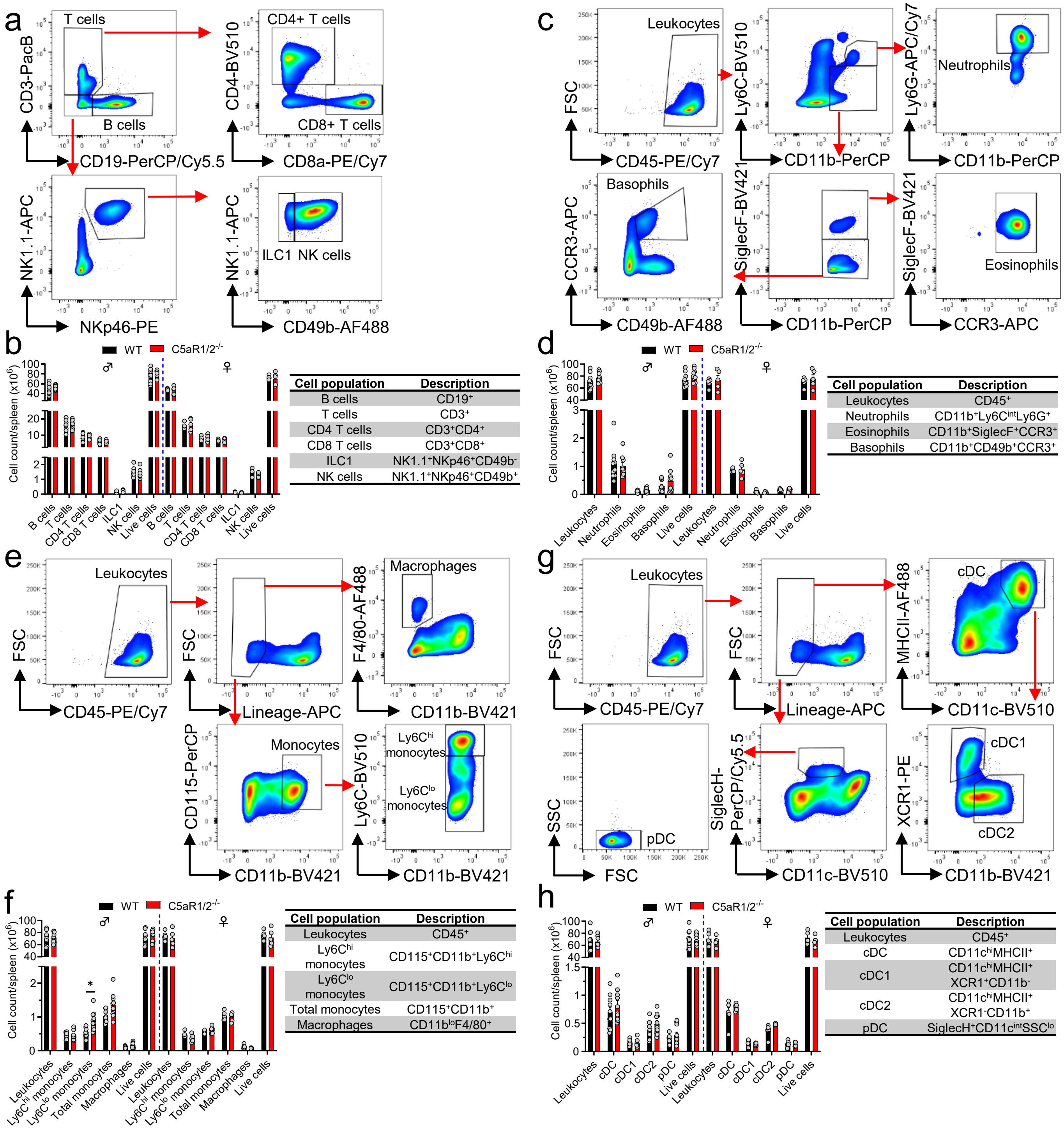


**Extended Data Fig.2: Immunophenotyping analysis of splenocytes from C5aR1/2$^{-/-}$ double knockout mice.** Related to Fig. 1. C5aR1/2$^{-/-}$ mice (females: ♀; males: ♂) and their C57BL/6J (WT) counterparts were necropsied at 17 weeks of age and spleens were collected to isolate immune cells by enzymatic digestion and analyzed by flow cytometry. (**a, c, e, g**) Representative gating strategy for splenic immune cells in naïve mice. After doublets and dead cells exclusion (eF780-FVD$^+$ cells), further gating was used to identify, (**a**) Lymphocytes (T cells, CD8$^+$ and CD4$^+$ T cells, B cells, NK cells, ILC1 cells), (**c**) Leukocytes (CD45$^+$ cells), Granulocytes (Neutrophils, Eosinophils, Basophils), (**e**) Monocytes/Macrophages, and (**g**) Dendritic cells (pDC, cDC, cDC1, cDC2). Lineage: CD3/CD19/CD49b/Ly6G. (**b, d, f, h**) Graphs show total immune cell counts in each spleen. Bars show mean ± SEM values from the analysis of n ≥ 5 mice per strain and per gender. Data are representative of 3 independent experiments. *$p < 0.05$ as determined by Student's t-test analysis. FVD, Fixable Viability Dye; NK, Natural Killer; ILC1, Innate Lymphoid cells 1; pDC, plasmacytoid DC; cDC, conventional DC; lo, low; int, intermediate; hi, high.

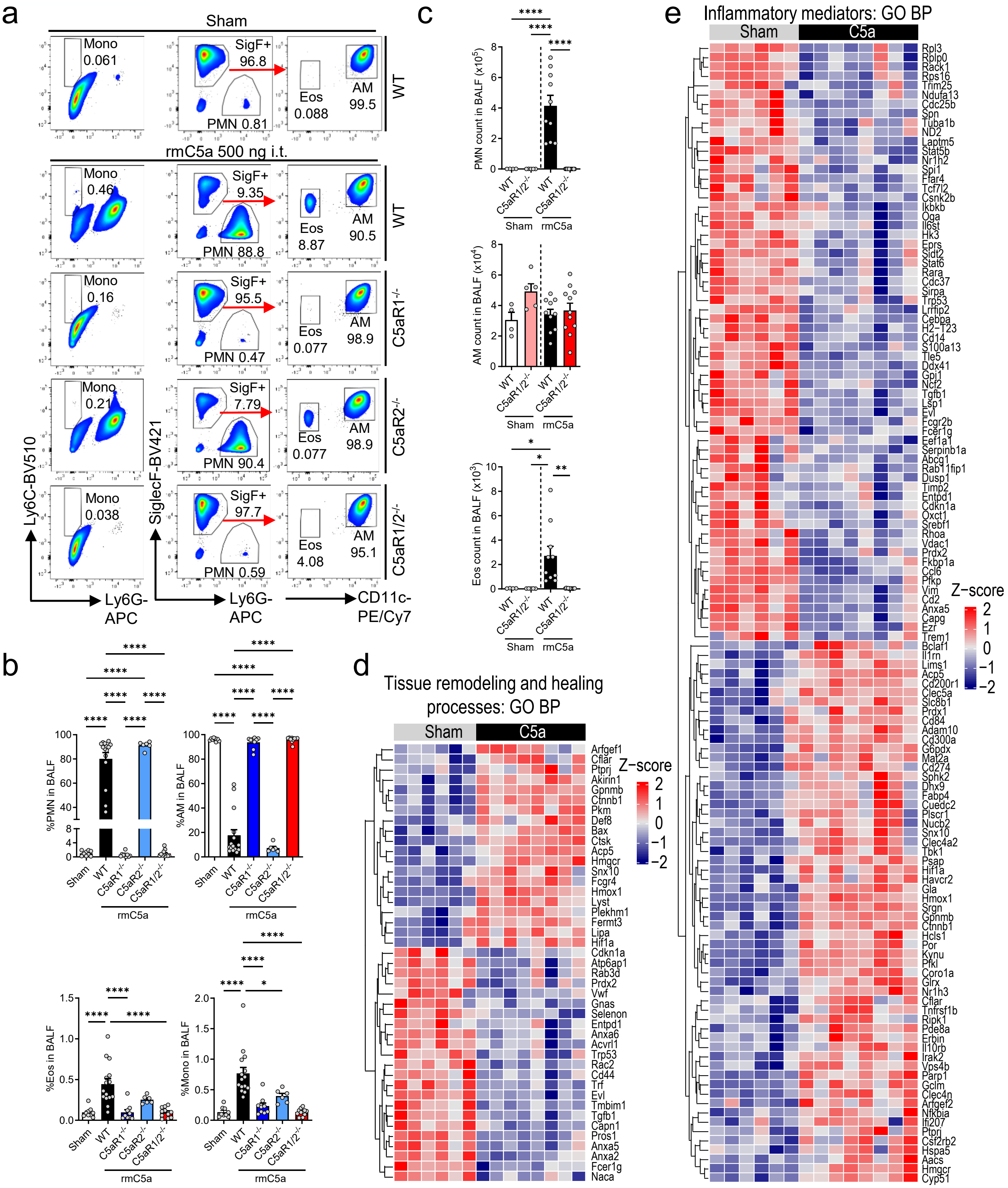


**Extended Data Fig.3: Transcriptional regulation of inflammatory mediators and genes involved in tissue remodeling and healing processes in alveolar macrophages (AM) following C5a-ALI in WT mice.** Related to Fig. 1 and 2. (**a**) Representative gating strategy, (**b**) cell frequencies and (**c**) cell counts of BALF infiltrating immune cell populations 7 h after rmC5a (500 ng) or PBS (sham) i.t. instillation in WT, C5aR1$^{-/-}$, C5aR2$^{-/-}$ and C5aR1/2$^{-/-}$ mice identifying Ly6G$^{+}$Ly6C$^{+}$ neutrophils (PMN), SiglecF$^{+}$CD11c$^{+}$ alveolar macrophages (AM), Ly6G$^{-}$Ly6C$^{+}$ monocytes (Mono) and SiglecF$^{+}$CD11c$^{-}$ eosinophils (Eos). Inset numbers indicate percentages of cells from parental gates falling into daughter gates. (**b-c**) Data are shown as mean ± SEM and are representative of 2-3 independent experiments. Each circle represents an individual mouse; (**b**) n = 6-15 mice/group, (**c**) n = 4-11 mice/group. *p < 0.05, **p < 0.01, ***p < 0.001, ****p < 0.0001, ns, not significant, as determined by one-way ANOVA. rmC5a, recombinant mouse C5a; BALF, bronchoalveolar lavage fluid. (**d-e**) Bulk RNA-seq was performed on FACS-sorted live SiglecF$^{+}$CD11c$^{+}$CD64$^{+}$ AM from sham and C5a-ALI mice (n = 6-8 mice/group). Heatmaps showing significant DEGs enriched in gene ontology biological processes (GO BPs) associated with (**d**) tissue remodeling and wound healing; (**e**) production, regulation, and response to inflammatory mediators (cytokines and chemokines) and reactive oxygen species metabolism. Heatmaps visualize normalized expression values of DEGs with baseMean ≥ 1000 with Z-score from -2/blue to +2/red. SigF, SiglecF.

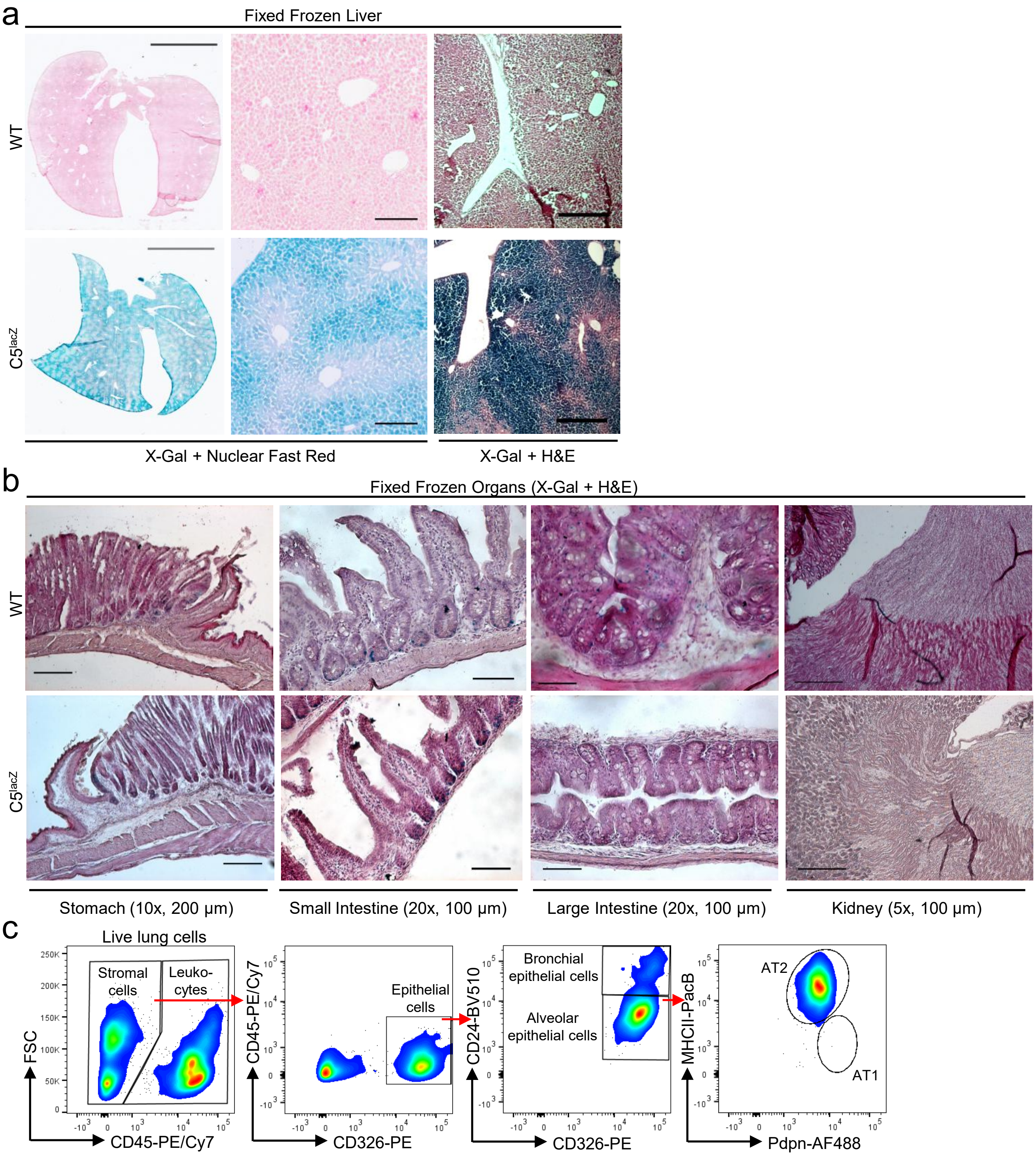


**Extended Data Fig.4: Identification of C5-producing cells in lungs of naïve reporter mice.** Related to Fig. 3. (**a-b**) Representative images of immunohistochemistry analysis for X-Gal and Nuclear Fast Red (**a**, left and middle panels) or H&E (**a**, right panels and **b**) staining of fixed-frozen liver (**a**) and indicated organs (**b**) sections from C5$^{lacZ}$ and WT control mice. (**a**, left and middle panels) Scale bars = 5 mm (whole section), 200 μm (zoomed sections). (**a**, right panel) Scale bars = 500 μm. (**b**) Scale bars are indicated under the respective panels. The data are representative of 3 experiments with similar results. (**c**) Dot plots showing the representative gating strategy used for alveolar type 2 epithelial cells (AT2) in enzymatically digested lungs of naïve C5$^{lacZ}$ mice by flow cytometry. Cell debris, doublets and dead cells were excluded prior to lung single-cell suspensions analysis. Pdpn, podoplanin.

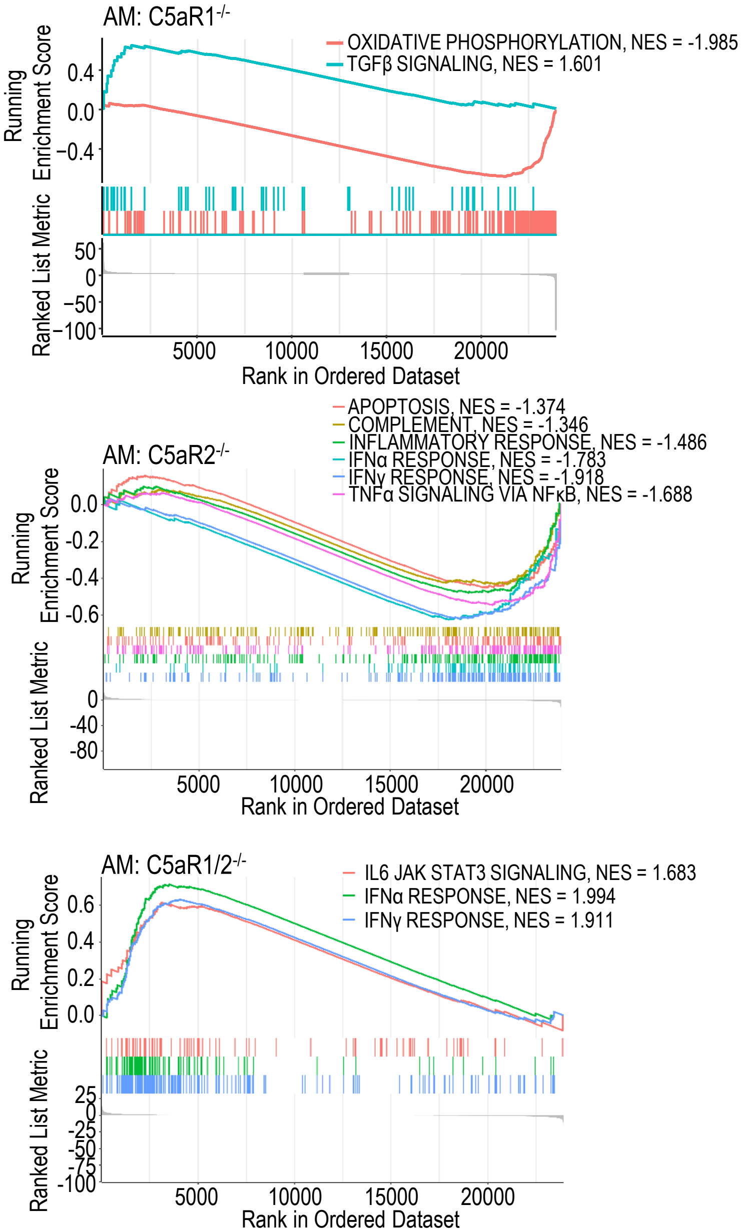


**Extended Data Fig.5: Loss of C5aR1 and C5aR2 differentially remodels inflammatory pathways in alveolar macrophages during *S. pneumoniae* infection in mice.** Related to Fig. 6. Bulk RNA-seq was performed on FACS-sorted live SiglecF$^+$CD11c$^+$CD64$^+$ alveolar macrophages (AM) from WT, C5aR1$^{-/-}$, C5aR2$^{-/-}$ and C5aR1/2$^{-/-}$ mice 18 h after intranasal infection with 2x10$^6$ CFU Spn4. Gene set enrichment analysis (GSEA) showing selected immune-related hallmark pathways significantly altered in C5aRs-deficient AM (adjusted $p < 0.05$) based on the analysis of significant and non-significant differentially expressed genes (DEGs). NES, normalized enrichment score.

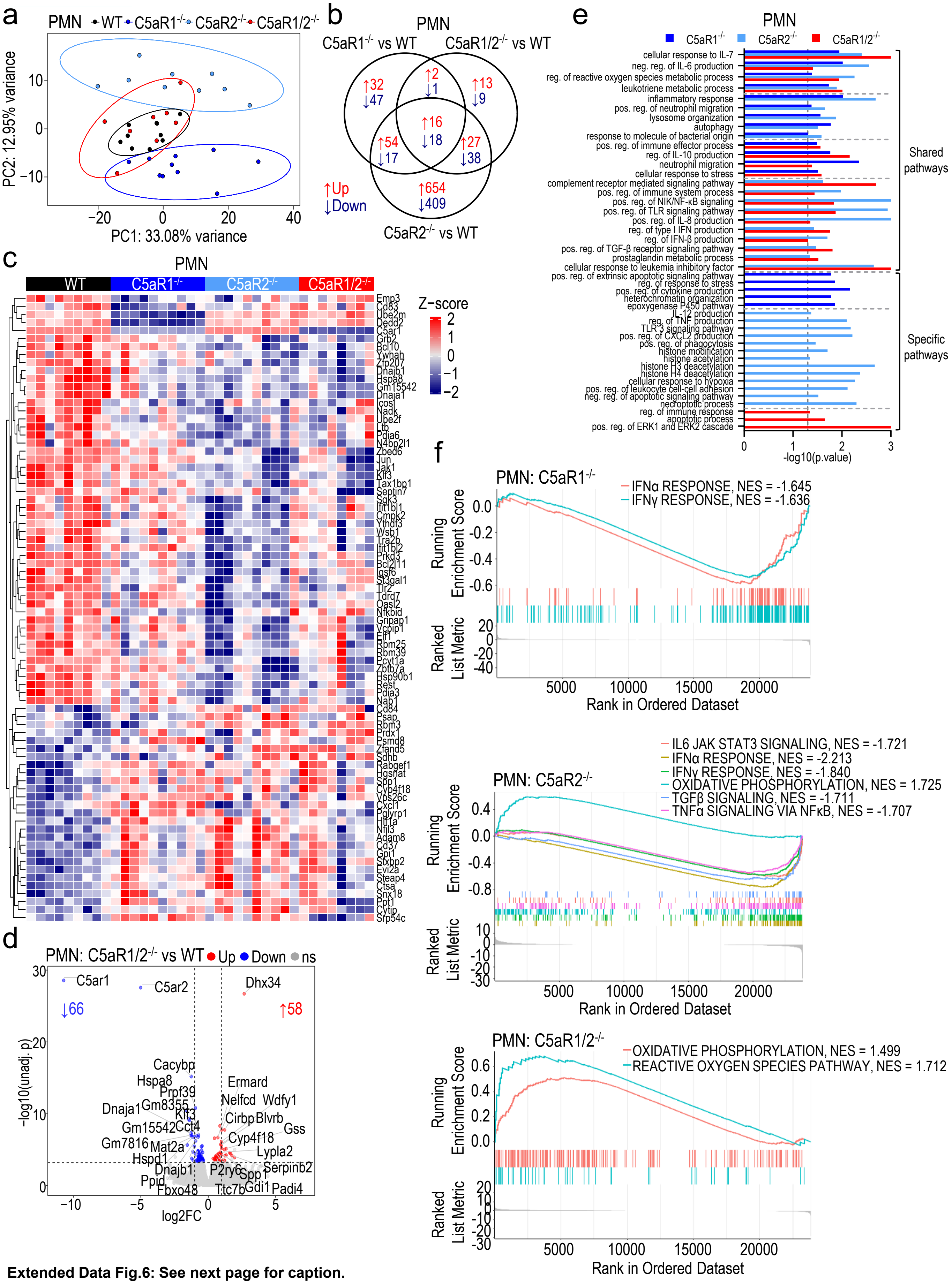


**Extended Data Fig.6: See next page for caption.**

**Extended Data Fig. 6: Single or combined deletion of C5aR1 and C5aR2 impaired neutrophil immune responses during pneumococcal infection in mice.**

Related to Fig. 6. Bulk RNA-seq was performed on FACS-sorted live Ly6G$^+$ alveolar neutrophils (PMN) from WT, C5aR1$^{-/-}$, C5aR2$^{-/-}$ and C5aR1/2$^{-/-}$ mice 18 h after intranasal infection with 2x10$^6$ CFU Spn4. (**a**) Principal component analysis (PCA) of the top 500 most variable genes after variance stabilizing transformation of RNA-seq counts (n = 8-10 mice/group). (**b**) Venn diagram showing shared and genotype-specific differentially expressed genes (DEGs). (**c**) Heatmap of normalized expression values of top 50 significant DEGs (baseMean ≥ 1000; Z-score from -2, blue to +2, red). (**d**) Volcano plot showing DEGs in C5aR1/2$^{-/-}$ vs. WT PMN. The number of significantly upregulated and downregulated genes is indicated, with top 15 genes with |log2FC| > 0.58, and C5aR2, labelled. The horizontal dotted line represents adjusted p (adj. p) = 0.05 and vertical lines represent |log2FC| = 1. FC, Fold Change; unadj. p, unadjusted p; ns, not significant. (**e**) Gene ontology (GO) biological processes (BPs) enriched among significant DEGs, highlighting immune-related pathways. pos., positive; neg., negative; reg., regulation. (**f**) Gene set enrichment analysis (GSEA) showing selected immune-related hallmark pathways altered in C5aRs-deficient PMN based on the analysis of significant and non-significant DEGs with Benjamini-Hochberg multiple testing for enrichment. GSEA visualization for C5aR1$^{-/-}$ vs WT PMN depicts processes with adjusted p < 0.2, although adjusted p < 0.05 was used to plot GSEA pathways for C5aR2$^{-/-}$ or C5aR1/2$^{-/-}$ vs WT PMN. NES, normalized enrichment score.

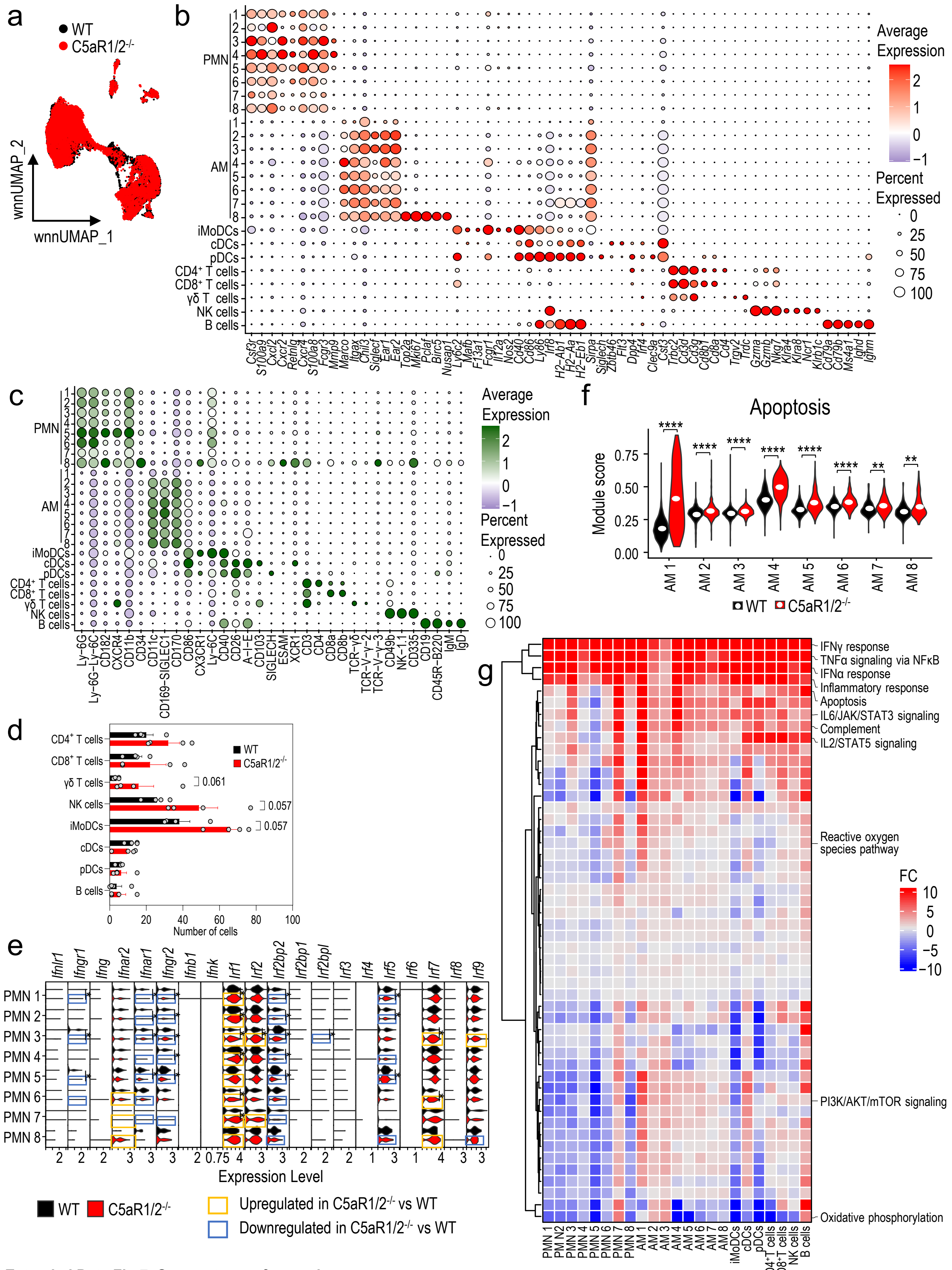


**Extended Data Fig.7: See next page for caption.**

**Extended Data Fig. 7: Single-cell sequencing of BALF leukocytes from WT and C5aR1/2$^{-/-}$ mice.**

Related to Fig. 7. Multimodal sequencing was performed using Cellular Indexing of Transcriptomes and Epitopes by Sequencing (CITE-seq) of FACS-sorted CD45$^{+}$ live BALF cells from WT and C5aR1/2$^{-/-}$ mice (n = 4/group) intranasally infected with 2x10$^{6}$ CFU Spn4 for 18 h. (**a**) Plot showing integrated weighted nearest neighbor uniform manifold approximation and projection (wnnUMAP) overlap in WT and C5aR1/2$^{-/-}$ mice. (**b-c**) Dot plot showing the expression of (**b**) gene and (**c**) protein markers used for cell type annotation. Panel **c** is an expanded plot from **Fig. 7b**. (**d**) Bar plot showing cell type distribution of T, NK, dendritic (DCs), B, and inflammatory monocytes/DC (iMoDCs) cells. pDCs, plasmacytoid DCs; cDCs, classical DCs. (**e**) Violin plot showing expression of interferon genes. (**f**) Violin plot showing module score based on apoptosis related GO BPs across AM clusters. (**g**) Heatmap showing fold change (FC) in multivariate distribution of hallmark gene sets between C5aR1/2$^{-/-}$ and WT mice. Higher/red indicates increase in gene set distribution in C5aR1/2$^{-/-}$ vs. WT. Selected immune-associated gene sets are labelled. *adjusted p < 0.05, **p < 0.01, ***p < 0.001, ****p < 0.0001.

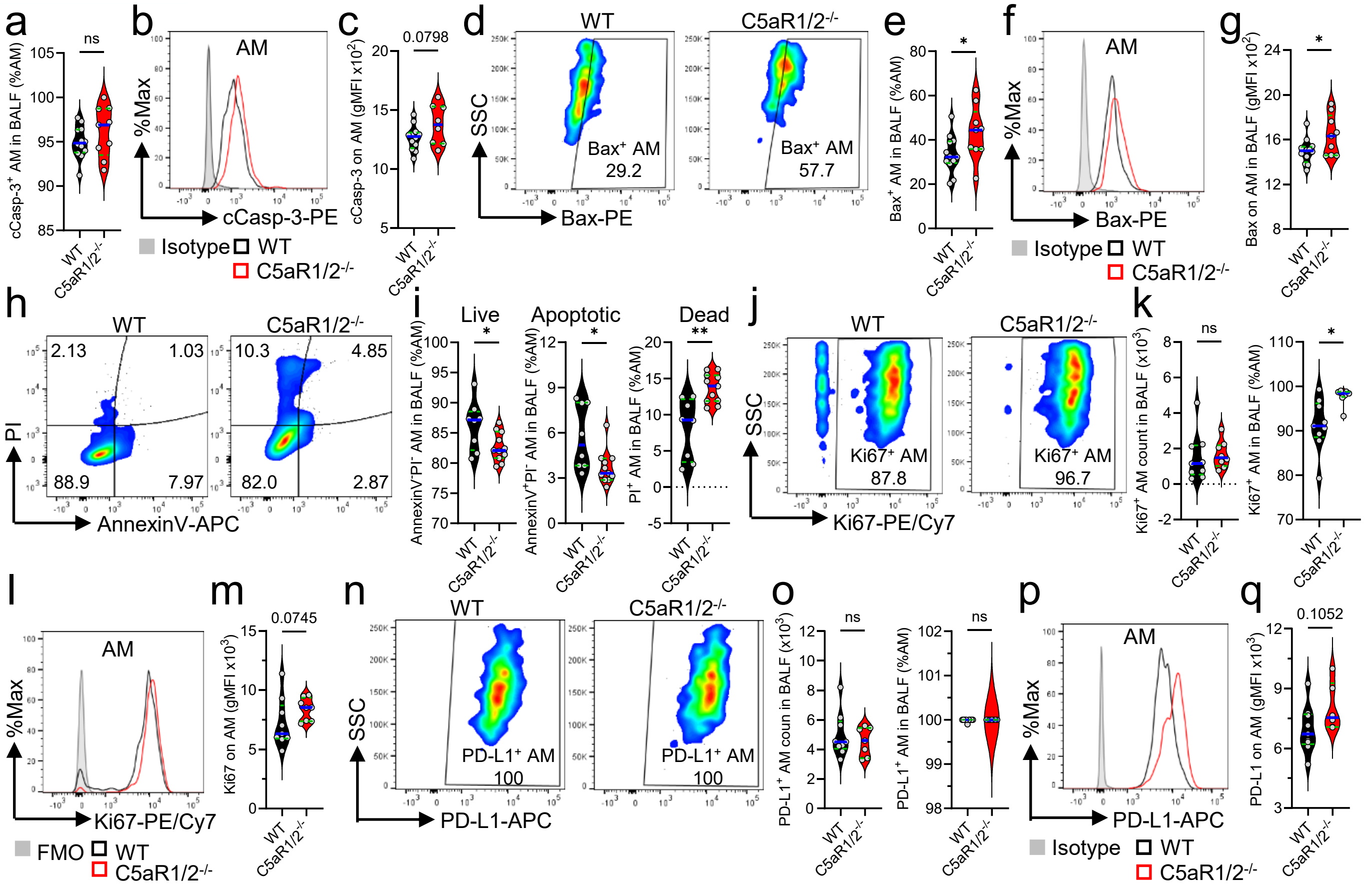


**Extended Data Fig.8: C5aR1/2$^{-/-}$ mice show differential alteration of AM proliferation and cell death during pneumococcal pneumonia.** Related to Fig. 8. WT and C5aR1/2$^{-/-}$ mice were intranasally infected with Spn4 (1-2x10$^6$ CFU/mouse) or PBS (sham), and BALF were collected 18 h later. (**a, e**) Percentages of live c-Casp-3$^+$ or Bax$^+$ total Siglec$^+$CD11c$^+$ alveolar macrophages (AM) were determined by flow cytometry (n ≥ 9/group). (**b, f**) Representative overlay histograms, and (**c, g**) gMFI of cCasp-3 or Bax on AM. Isotype control was included as a reference (n ≥ 8/group). (**d**) Representative flow cytometry plots of Bax on AM from BAL. Cell frequencies of total AM are indicated next to each gate. (**h**) Flow cytometric analysis measuring apoptosis in AM using AnnexinV and Propidium Iodide (PI) staining. Cell frequencies of total AM are indicated next to each gate. (**i**) Bar charts show the percentage of live (AnnexinV$^-$PI$^-$), apoptotic (AnnexinV$^+$PI$^-$) and dead (PI$^+$) AM in both groups (n ≥ 8/group). (**j**) Representative flow cytometry plots and, (**k**) cell count and fraction of AM that were actively proliferating, expressed as the percentage of total Ki67$^+$ AM (n = 8-9/group). (**l**) Representative overlay histograms and (**m**) gMFI of Ki67 on AM. FMO was included as a control (n = 8-9/group). (**n**) Representative flow cytometry dot plots, and (**o**) cell count and fraction of total AM expressing PD-L1 (n = 7-9/group). (**p**) Representative overlay histograms and (**q**) gMFI of PD-L1 expression on PMN. Data are shown as mean ± SEM and are representative of 2-3 independent experiments with similar results. Each circle represent an individual mouse. *p < 0.05, **p < 0.01, ***p < 0.001, ****p < 0.0001, as determined by Student's t-test. BALF, bronchoalveolar lavage fluid; FMO, fluorescence Minus One; gMFI, geometric mean fluorescence intensity.
